# KLHDC7B, a novel gene associated with age-related hearing loss in humans, is required for the maintenance of hearing in mice

**DOI:** 10.1101/2024.10.14.618265

**Authors:** Alexandra M. Kaufman, Benjamin Silver, Roberto A Donnianni, Carlos Aguilar, Lingzi Niu, Daniel Johnson, Anwen Bullen, Alma Corona, Benjamin J. van Soldt, Nilay Vora, Gervasio Batista, Luz Cortes-Burgos, Jacqueline Copeland, Elika Fallah, Norman Zhang, Marina Lehmkuhl, Sarah Cancelarich, Kara Campos, Daniela Di Battista Miani, Jaylen Mumphrey, Susan D Croll, Johnathon R. Walls, Mary Germino, Michael R Bowl, Meghan C Drummond, Sally J. Dawson

## Abstract

**Background:** Age-related hearing loss (ARHL) is the most common sensory loss in older adults, but the underlying pathological mechanisms remain unclear. Recent genome wide association studies (GWAS) have linked variation in a large number of genes with increased risk of ARHL for the first time. Amongst the strongest of these associations is variation in *KLHDC7B,* a gene of unknown function and one not previously linked to hearing.

**Methods:** To confirm whether KLHDC7B plays a role in hearing we investigated auditory function in two independent knockout mutant mouse models of *Klhdc7b: Klhdc7b^IMPC−/−^*and *Klhdc7b^RegnΔ/Δ^* on the C57BL/6N background and the B6.CAST-*Cdh23^753A>G^* background respectively. The B6.CAST-*Cdh23^753A>G^* background was backcrossed to correct a known age-related hearing loss (*ahl*) mutation in *cadherin 23* present in both the C57BL/6N and C57BL/6J strains.

**Results:** We showed that *Klhdc7b* is expressed exclusively in inner and outer sensory hair cells within the cochlea of mice at the RNA and protein level. Homozygous mutants for both knockout mouse models display a similar early-onset, progressive and severe hearing loss. Histological characterization of the two mouse models suggests that hair cells develop normally and are present in neonates. However, after the onset of hearing there is a progressive loss of outer hair cells in a gradient from base to apex of the cochlea consistent with the hearing deficit in the mice and the pattern of hearing loss in ARHL. Inner hair cells remain intact up to the latest age examined (~8 weeks).

**Conclusions:** Our data suggests KLHDC7B is required for maintenance of auditory function rather than in its development, supporting the novel association with ARHL in humans detected in recent GWAS. To our knowledge, this is the first validation in mouse of an ARHL association in humans detected in a GWAS. Our work also provides two distinct mouse models to further investigate the role of KLHDC7B in auditory function and for use in the development of therapeutic tools to prevent or treat ARHL.

## Background

Age-related hearing loss (ARHL) is a common condition that can lead to social isolation, depression and a decrease in the quality of life. Furthermore, hearing loss in mid-life has recently been found to be associated with an increased risk of subsequent dementia diagnosis. In 2020, the Lancet Commission published an update on their previous report in 2017 which again pinpointed hearing loss in mid-life as the largest modifiable risk factor for dementia (1, 2). The number of older people with hearing loss and dementia is rising as the aging population increases (GBD 2019 Dementia Forecasting 3). Age-related hearing loss is the most common sensory loss with more than 50% of people aged 60-69, and over 80% of those aged 85 years and older having some hearing loss that affects daily communication (4). It is a complex condition involving a combination of genetic risk factors and exposure to environmental components such as noise exposure and ototoxic drugs. Evidence from early GWAS into ARHL suggested that genetic effects were likely to be small or due to rare alleles, and these studies largely did not identify genome-wide significant associations (5–12). However, several recent GWAS utilizing self-report hearing data in the large UK Biobank Cohort dataset, together with subsequent meta-analysis and replication studies have for the first time identified over 50 genetic loci significantly associated with ARHL risk (13–17). Some of these genes have known auditory function and/or have been previously linked to congenital deafness but the majority are novel associations with hearing thereby presenting an opportunity to reveal new pathogenic mechanisms for ARHL.

The strongest association (p=1.90E-22) with self-reported hearing in the UK Biobank GWAS (Wells et al., 2019) was with a Val504Met missense variant (rs36062310) within the kelch-domain containing 7b (*KLHDC7B*) gene. A recent meta-analysis of five cohorts identified the same variant in association with hearing loss in adults (p=4.24E-26) (17). The same study also utilized whole exome sequencing data to perform a rare variant analysis (minor allele frequency <0.01), identifying heterozygous predicted loss-of-function variants in *KLHDC7B* associated with increased risk of hearing loss, with an odds ratio of 2.145. Taken together these genetic associations provide a strong rationale for *KLHDC7B* being a novel candidate gene for ARHL, but KLHDC7B has not yet been studied in the auditory system.

Other than its membership in the Kelch-domain containing protein superfamily (due to the presence of highly conserved Kelch domains), very little is known about this protein. Kelch domains are a set of repeating beta-sheet forming subunits that come together to form a tertiary structure known as a beta propeller. Kelch domain containing proteins have diverse subcellular locations and functions (18, 19), and there are only a few publications specifically examining KLHDC7B. It has been shown to be upregulated, yet also hypermethylated, in breast cancer cells (20), but its biological function remains uncharacterized.

To investigate the role of *KLHDC7B* in hearing, we determined the cell-type specific expression of *Klhdc7b* in the mouse cochlea using qPCR, RNAscope and immunofluorescence finding that within the cochlea, *Klhdc7b* is exclusively detected in hair cells, including inner, outer, and vestibular hair cells. To determine whether KLHDC7B is required for auditory function we utilized two independent knockout mouse models, *Klhdc7b^IMPC−/−^* and *Klhdc7b^RegnΔ/Δ^.* The first was on a C57BL/6N background and the second was on the B6.CAST-*Cdh23^753A>G^* background, a line that was bred with the CAST/EiJ line and then backcrossed to C57BL/6J to produce a C57BL/6J line carrying the wildtype allele of *Cdh23* rather than the known age related hearing loss allele in *Cdh23* (21).

We performed auditory phenotyping by auditory brainstem response (ABR) of wildtype, heterozygous and homozygote mutant mice in both models at several ages. Mice homozygous for either knockout allele display a similar pattern of severe early hearing loss with some residual hearing progressing to profound deafness at 8-12 weeks of age. Distortion product otoacoustic emission (DPOAE) measurements indicate a similar progression of loss of outer hair cell function in *Klhdc7b^RegnΔ/Δ^* and *Klhdc7b^IMPC−/−^* mice. Histological assessment of cochlear whole mount preparations, scanning electron microscopy (SEM) and transmission electron microscopy (TEM) from these mice indicate that outer hair cells develop normally but begin to die at around the same time as onset of hearing, around two weeks after birth. Taken together, our results confirm KLHDC7B is required for auditory function and suggest a role in outer hair cell maintenance beginning at the onset of hearing, a role that is consistent with its implication as an ARHL susceptibility gene.

## Methods

### Expression Probe Design and qPCR

qPCR probes were designed using BioSearch RealTimeDesign qPCR Assay Design software (https://www.biosearchtech.com/support/tools/design-software/realtimedesign-software) and tested for specificity via the UCSC genome browser BLAT function. The mouse L+S *Klhdc7b* probe was designed to a portion of the transcript that overlaps the two putative long and short isoforms. The mouse L *Klhdc7b* probe was designed against a portion of the transcript that is only present in the long isoform. For sequences, see supplemental methods. For tissue collected in-house, mice under 7 days old were decapitated; older mice were euthanized under CO_2_ with secondary decapitation. Organs were placed in RNAlater®, cochleae were frozen on dry ice. For RNA extractions, tissue/cells were homogenized in TRIzol®, and chloroform was used for phase separation. Total RNA was purified using MagMAX™-96 for Microarrays Total RNA Isolation Kit (Ambion by Life Technologies) according to manufacturer’s specifications. Genomic DNA was removed using RNase-Free DNase Set (Qiagen). mRNA was reverse-transcribed into cDNA using SuperScript® VILO™ Master Mix (Invitrogen by Life Technologies) and amplified with the SensiFAST Probe Lo-ROX (Meridian) using the 12K Flex System (Applied Biosystems).

### Animal husbandry

All *Regn* mice were housed and treated according to guidelines from the Institutional Animal Care and Use Committee. All mice used for hearing experiments were crossed and bred to homozygosity with the B6.CAST-*Cdh23^753A>G^* line (21). Both male and female mice were used. All *IMPC* mice were housed in individually ventilated cages with environmental conditions according to UK Home Office regulations. All procedures were licensed by the Home Office under the Animals (Scientific Procedures) Act 1986 and Amendment Regulations 2012 (Project Licence numbers PP7565374 and PP8324029) and approved by local institutional ethical review committees. All mice were culled using methods approved under these licences to minimise any possibility of suffering. Both male and female mice were used.

### Knockout mouse generation

For *Klhdc7b^RegnΔ/Δ^* mice, a targeting vector for knocking out (KO) an endogenous *Klhdc7b* gene was constructed using bacterial artificial chromosome (BAC) clones and VELOCIGENE® technology (see U.S. Patent No. 6,586,251) (22). For details, see Supplemental Methods. The modified BAC clone containing the KO of *Klhdc7b* gene, LacZ and a self-deleting Neomycin cassette was used to electroporate mouse embryonic (ES) cells to create modified ES cells comprising a *Klhdc7b* KO gene into B6 mice and content confirmed by TaqMan assay. Female mice were implanted using the VELOCIMOUSE® method (see, e.g., U.S. Pat. No. 7,294,754). Briefly, ES cells were electroporated into an early-stage blastocyst (8 cells Morula stage) to generate F0 mice that are essentially fully derived from the donor gene-targeted ES cells allowing immediate phenotypic analyses (23). The Neomycin selection cassette was removed by crossing the progeny generated from the ES clone with a deletor rodent strain that expresses a Cre recombinase. Animals homozygous KO for the *Klhdc7b* locus are bred by crossing heterozygous animals. For sequence descriptions, primers and probes used, see supplemental methods.

The ***H-KLHDC7B-DEL1258-EM1-B6N*** mouse line (designated here as *Klhdc7b*^IMPC*−/−*^) was generated at MRC Harwell as part of the IMPC program (24) via microinjection of CRISPR/Cas9 reagents into primed C57BL/6N oocytes which were implanted in pseudo-pregnant CD1 surrogate females. The CRISPR-induced mutation created a deletion of 1258bp in the single exon of the *Klhdc7b* gene, within the sequence common to both long and short isoforms of *Klhdc7b*. In addition to removing 419 amino acids, the deletion also induces a premature stop codon creating a null allele (for further details see https://www.mousephenotype.org/data/genes/MGI:3648212). Four *IMPC^+/−^* males and four *IMPC^+/−^* females were imported from the MRC Mary Lyon Centre, MRC Harwell and a colony was maintained at UCL. Occasionally, *IMPC^+/−^* mice were outcrossed to C57BL/6N mice with the aim to offset allelic drift. Mice were genotyped by rt-PCR on cDNA made from ear biopsies to detect wildtype and mutant alleles – for primers, see Supplemental Methods.

### Histology: tissue processing and staining

#### Animal sacrifice and cochlea treatment

For Regeneron mice, animals 21 days and older were sacrificed by transcardial perfusion with phosphate buffered saline (PBS) followed by 4% paraformaldehyde (PFA) in PBS. The cochleae were extracted from the temporal bone. A tiny hand-drilled hole was made on the apex, the stapes was removed and oval window pierced. Cochleae were placed 1-4 hours or overnight in 4% PFA followed by 3x PBS washes and stored at 4°C. The tissue was decalcified in Immunocal® (Statbio) overnight and rinsed 3x with PBS.

For UCL *Klhdc7b^IMPC^* mice, animals were sacrificed according to Schedule 1 procedures as described in the United Kingdom (Scientific Procedures) Act of 1986. For preparation of cochlea, whole inner ear was dissected (Montgomery and Cox, 2016) and fixed in 4% paraformaldehyde (PFA) diluted in 10 mM phosphate buffered saline (PBS) pH 7.4 for two hours. Samples with age above P4 were then decalcified in 4% Ethylenediaminetetraacetic acid (EDTA) for 48 hours.

#### Whole mount immunostaining

For immunohistochemistry of whole-mounted Organ of Corti, the sensory epithelium was carefully dissected after decalcification. One of two protocols were used. For *Klhdc7b^IMPC^* data, tissue was permeabilized with 5% tween-20 in PBS for 1 h at room temperature and blocked with 10% goat serum, 0.5% Triton-X 100 in PBS for 2 h at room temperature. Incubation with primary antibodies was performed overnight at 4°C. After three washes using PBS for 15 min, the secondary antibody incubations were undertaken of 1 h at room temperature in the dark. Nuclei were visualized with 1µM DAPI, 10nM phalloidin-Atto 647N (Sigma-Aldrich, Gillingham, UK).

For *Klhd7cb^Regn^* data, dissected samples were washed 3x in PBS, incubated with blocking solution for 1 hour at room temperature, incubated overnight with primary antibodies in blocking solution at 4°C or room temperature, then washed 3 x with PBS. Secondary antibodies and cell stains were diluted in blocking buffer and samples were incubated for 1 hour at room temperature, rinsed 3x with PBS, then placed on slides and covered with Prolong gold and a coverslip. For antibodies and concentrations, see supplemental methods.

#### Paraffin embedding, slicing and immunostaining

For paraffin embedding, cochleae were placed in cassettes, dehydrated and immersed in a vacuum chamber containing warm paraffin wax using a tissue processor (Leica). The protocol was 70% ethanol for 10 min, 95% ethanol for 15 min, 95% ethanol for 12 min, 100% ethanol for 12 min x 3, xylene x 12 min, xylene x 15 min x 2, and paraffin x 30 min, then 35 min. Cochleae were embedded in paraffin, sliced using a microtome at six-micron thickness and dried on a heat block. For immunostaining paraffin-embedded sliced cochlea, slides were baked at 60°C for one hour, washed in xylene 2x for 3 minutes. Slides were rehydrated in 100% ethanol 2x 3 minutes, then 95% ethanol 2x 3 minutes, then 70% ethanol for 3 minutes, 50% ethanol for 3 minutes, distilled water for 5 minutes, and 3x 2 minutes PBS. Antigen retrieval was performed using Co-detection target retrieval reagent (ACD Bio). This was heated in a vegetable steamer (Oster), and slides were placed in hot co-detection target retrieval reagent for 20 minutes in the steamer, then allowed to cool for 10 minutes, followed by two washes PBS plus 0.5% Tween-20.

A hydrophobic barrier was drawn around the sections and slides were placed in a humidified chamber and covered with blocking solution (2%w/v bovine serum albumin, 5% normal donkey serum, 0.01 % triton-x 100 in PBS), then incubated at room temperature for 30 min. Primary antibodies diluted in blocking solution were added to the slides and incubated at 4°C overnight. Slides were washed 3x with PBS, then covered with secondary antibodies and cell stains diluted in blocking solution and incubated for 1 hour. Afterward, slides were washed 3 x in PBS, then mounted using Prolong Diamond or Prolong gold and covered with a coverslip.

#### Cryoembedding and staining

For cryoembedding and sectioning, decalcified cochleae were dehydrated in 15% and 30% sucrose successively and embedded in optimal cutting temperature media in liquid nitrogen. Cryoembedded samples were serially sectioned in a 10 µm thickness using a OTF6000 Cryostat (Bright Instruments, UK). Mid-modiolar sections was collected for subsequent immunostaining. Cryosections were initially rehydrated in PBS for 10 min and incubated in 1% sodium dodecyl sulfate (SDS) for 5 min to retrieve antigen at room temperature. After three washes in PBS for 10 min each time, sections were permeabilized and blocked in blocking buffer composed of 10% goat serum, 0.1% Triton-X100 in PBS for 1 hour at room temperature. Sections were then incubated with primary antibodies overnight at 4°C, followed by three washes with PBS-T (0.1% Triton-X100 in PBS) for 10 min each. Secondary antibodies and DAPI were diluted in blocking buffer, in which sections were incubated for 2 hours at room temperature. After three washes with PBS-T, sections were mounted in Fluoromount-G® (SouthernBiotech).

#### RNAscope with antibody co-detection

For RNA scope in paraffin embedded sections, the ACD Bio protocol for paraffin embedded slides was followed using the suggested standard incubation times for antigen retrieval. Antibody (MYO7A, Proteus, 25-6790) was diluted at a concentration of 1:200 in Co-detection antibody diluent and incubated overnight at 4°C in a humidified chamber. Opal 520 and 570 dyes were used at 1:1500. For details on reagents, see supplemental methods.

#### LacZ staining of knockout mice

Mice were anesthetized with ketamine/xylazine and fixed via perfusion with 2% paraformaldehyde. After perfusion, tissues were dissected and sliced into 1-5 mm pieces, fixed 30 min at room temperature, washed in PBS for 30 min and stained in beta galactosidase (LacZ) staining solution overnight at 4°C. After staining, tissues were washed in cold PBS for 15 min and postfixed in 4% formaldehyde at 4°C overnight with mixing. Tissues were cleared with glycerol by incubating in 50% glycerol for a day at RT and then 70% glycerol for a day. Tissues were photographed on a Zeiss dissection microscope and stored at RT in 70% glycerol. Tissues were subsequently decalcified overnight with Immunocal® and dissected, then photographed using an Axioscan slide scanner (Zeiss).

#### Klhdc7b^IMPC^ hair cell counts

Hair cell counts were performed on maximum intensity projections of confocal Z-stacks collected at 1.0 µm intervals (63x oil, 1.4 N.A. objective) from a minimum of 3 mice for each genotype and age. The Z position was programmed consistently for each image stack between the highest point of the first stereocilia bundle to the lowest synaptic marker of the last IHC. Data were collected from mid-basal, mid-middle and mid-apical cochlear turns and correlated to the mouse tonotopic frequency-place map (25) for consistency. Data were quantified in 250 µm lengths and individual hair cells were counted on 3 separate occasions to ensure consistency.

### Confocal Microscopy

Samples were imaged on a confocal microscope; either Zeiss LSM 780 or Zeiss LSM 880 using 10x (0.45 N.A.), 20x (0.8 N.A.), and 63x oil (1.4 N.A.) objectives and Zen 3.0 SR software (Black edition). For figure 1D, Zeiss LSM 980 with Airyscan acquisition and processing were used, using Zen Blue software. Z-stacks were taken with the appropriate size for the objective and numerical aperture. Post hoc image processing and analysis was performed with Fiji. Saturation of images and zero black levels were avoided in image capture. For display, any adjustments were made equally to images from all 3 genotypes. All immunostaining experiments used a minimum of 3 mice per genotype and age range, representative images are shown in the figures.

**Figure 1.**
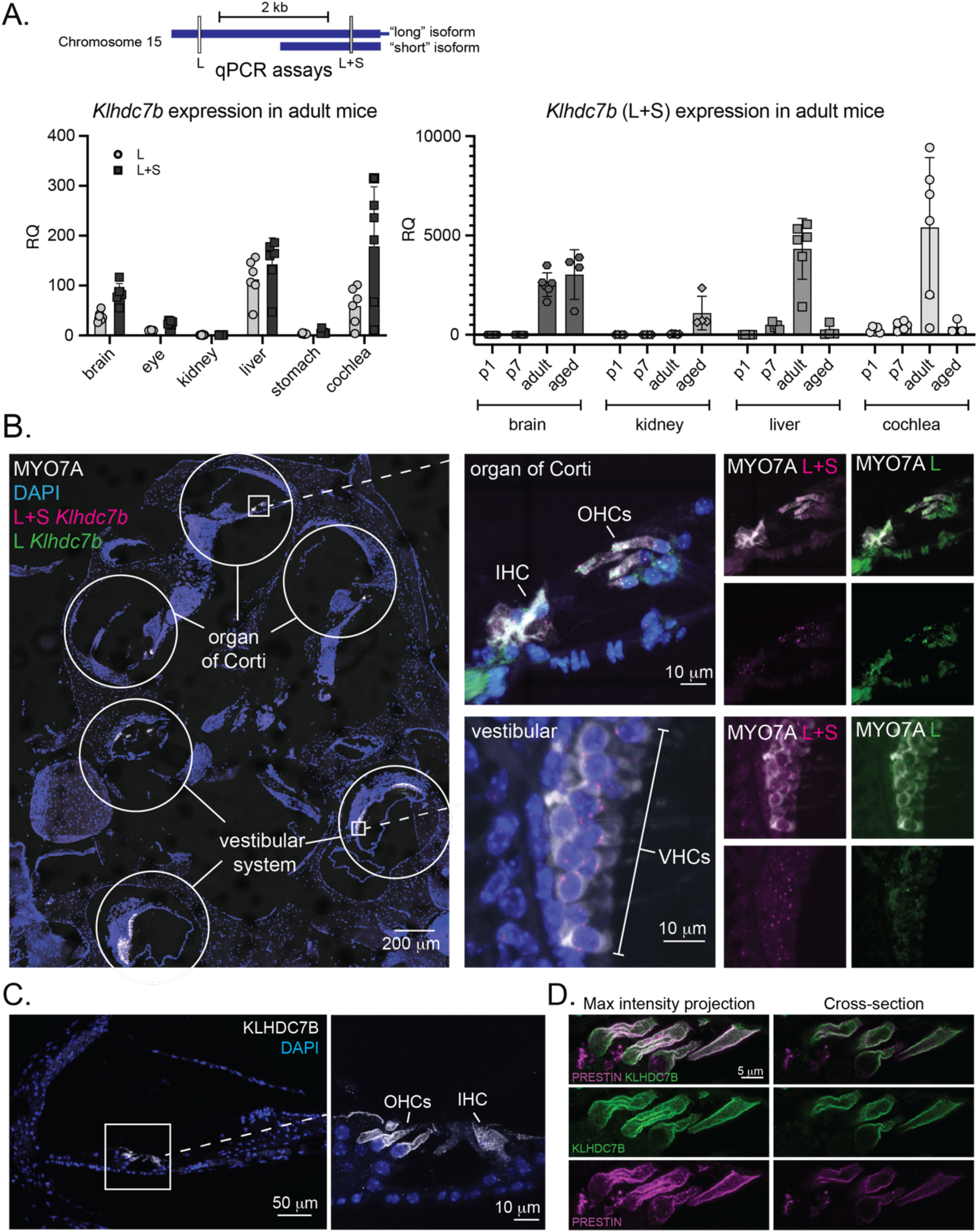
*Klhdc7b* is expressed in cochlear hair cells. **A.** qPCR primer/probe sets were designed against the mouse long isoform *(L)* and overlapping portions of the long and short (*L+S*) isoforms (schematic, top left). Left chart: qPCR using *Klhdc7b L* and *L+S* probes in cDNA derived from various adult mice tissue. RQ for each sample is normalized to *Drosha*, a housekeeping gene, and shown compared to mean of adult kidney as a calibrator. Right chart: qPCR of L+S isoform expression of cDNA derived from mouse tissues at different ages. RQ is normalized to *Drosha*, a housekeeping gene, then compared to mean of P1 kidney. P1, postnatal day 1. P7, postnatal day 7. P1 and P7. **B.** RNA scope probes were designed similarly to qPCR probes in **A** to recognize the overlapping and long portions of the transcripts. Left, RNAscope probes as indicated and MYO7A immunofluorescence (hair cell marker) and DAPI stain (nuclei) in adult cochlea. Top right, merged higher magnification confocal images of organ of Corti. Bottom right, merged higher magnification confocal images of vestibular hair cells. Far right, images of individual channels corresponding to organ of Corti and vestibular hair cells (merged MYO7A and L+S, MYO7A and L, with corresponding probe alone below). IHC, inner hair cell; OHC, outer hair cell; VHC, vestibular hair cell. **C.** One turn of adult organ of Corti showing immunofluorescence staining with an antibody against KLHDC7B. Left, 20x, right, 40x. **D.** Outer hair cells showing immunofluorescence for anti-KLHDC7B and anti-prestin, a protein localized to OHC membranes. Left, maximum intensity projection, right, single plane showing a cross-section of outer hair cells.

### Scanning Electron Microscopy (SEM) and Transmission Electron Microscopy (TEM)

For SEM of *Klhdc7b^Regn^* mice, after cochleae were removed from the temporal bone, they were fixed overnight in a mixture of 3% formaldehyde, 3% glutaraldehyde in 0.1 M sodium cacodylate buffer, pH 7.4 (Electron Microscopy Sciences, Hatfield, PA) supplemented with 2 mM of CaCl_2_. The samples were then washed with 0.1 M sodium cacodylate buffer, decalcified for 24 hours in Immunocal, washed 3X, and dissected to three flat turns. The bone, strial ligament and tectorial membrane were carefully removed from each turn and cochlear sections were placed into porous baskets. Afterwards, the samples were dehydrated through a graded series of ethanol (5%, 10%, 20%, 40%, 60%, 80% and 100%), critical-point dried using a manual CPD (BAL-TEC 030) in liquid CO_2_, sputter coated with 5nm Platinum and imaged with a field-emission scanning electron microscope (Zeiss Sigma VP).

For SEM of *Klhdc7b^IMPC^* mice, SEM preparation of the organ of Corti was performed as in previously (26). In brief, cochleae were fixed (2 h, 2.5% glutaraldehyde, 2% PFA 0.1 M cacodylate buffer, 3 mM CaCl_2_, room temperature) and decalcified (48 h, 4% EDTA, 4 °C) and the organ of Corti dissected, post-fixed in OsO_4_ and processed through the thiocarbohydrazide-Os-repeated procedure. Processed samples were dehydrated in a graded ethanol series, critical point-dried and sputter coated with platinum. Samples were examined in a JEOL 6700F SEM operating at 5 kV by secondary electron detection. Imaging was carried out using SEM Supporter software (System In Frontier, Japan).

For TEM, samples were prepared as described in Bullen, Forge (26). Briefly, samples were fixed and decalcified before post-fixation in OsO_4_. Cochlea was dehydrated through 30% and then 50% ethanol for 15 min each at room temperature. En-bloc staining was performed using 2% uranyl acetate in 70% ethanol at 4°C overnight. The next day, cochlea processed in 85%, 95%, 100% ethanol and propylene oxide for 15 min each, followed by incubation in 25% over-day, 50% overnight and 75% resin over-day. Cochleae were then embedded in 100% Agar100 Epon resin (Agar Scientific, UK) and let cure for 24 hours at 60°C. Resin blocks were trimmed and cut in a thickness of 110 nm using Ultramicrotome (Reichert). 100 nm sections were post-stained with uranyl acetate and lead citrate, before being examined on a JEOL 1400Flash transmission electron microscope operating at 120kV and equipped with a Gatan RIO16 camera.

### 3D microscopy

Cochleae were decalcified and carefully dissected to allow antibody penetration. Cochleae were stained with MYO7A and Sytox Deep red, a nuclear stain, embedded in agarose, and dehydrated using a series of increasing concentrations of methanol. Samples were cleared in ethyl cinnamate (ECi) and imaged using an Ultramicroscope Blaze (Miltenyi). For analysis, imaging files were converted to Imaris format, and hair cells were virtually dissected (27). Inner and outer hair cell masks were created, and data within the masks were exported to Python for further analysis. Hair cells were extracted using a custom cell segmentation algorithm. A custom processing pipeline was developed to unwrap the cochlea in a sequence that reflects its natural configuration and normalize it to the tonotopic map. For details, see supplemental methods.

### Behavioral assays

Mice were tested on open field, rotarod, pole test, and a Y-maze on different days. For details, see supplemental methods.

### Auditory brainstem response (ABR)

For *Klhdc7b^Regn^* mice, ABRs were performed in a soundproof booth (IAC Acoustics, IL). Recordings were performed at three pure-tone frequencies (8 kHz, 16 kHz, and 32 kHz) using the TDT RZ6 recording system (Tucker-Davis Technologies, FL), with 512 presentations of each stimulus averaged at each sound-pressure-level presented. Stimuli were presented for 2.5 ms with a 0.2 ms cos^2^ rise-decay with a hanning window applied at a rate of 21 presentations per second. Each frequency was played starting at a sound pressure level of 90 decibels (dB SPL), decreasing in 10 dB decrements to 50 dB and 5 dB decrements from 50 dB to 15 dB. The waveforms of the ABR were bandpass filtered between 300 Hz and 3 kHz with a 60 Hz notch filter. ABR recordings were processed in Matlab. Traces were first smoothed with a moving median filter with a kernel of 50 time points (each 10 ms recordings contained 244 time points). This was to remove slow wave noise occasionally observed in recordings and had a minimal effect on recordings that were not noisy. ABR thresholds were called manually by two experimenters on traces presented in a blinded, random order, and run through an algorithm adapted from the Liberman lab (28). See supplemental methods for more details.

For the *Klhdc7b^IMPC^* mice, ABR tests were performed at MRC Harwell using a click stimulus in addition to frequency-specific tone-burst stimuli to screen mice for auditory phenotypes and investigate auditory function. ABR responses were collected, amplified, and averaged using TDT System 3 (Tucker Davies Technology) in conjunction with BioSig RZ (v5.7.1) software. The TDT system click ABR stimuli comprised clicks of 0.1 ms broadband noise spanning approximately 2 - 48 kHz, presented at a rate of 21.1 s^−1^ with alternating polarity. Tone-burst stimuli were of 7 ms duration, inclusive of 1 ms rise/fall gating using a Cos^2^ filter, presented at a rate of 42.5 s^−1^ and were measured at 8, 16 and 32 kHz. All stimuli were presented free-field to the right ear of the mouse, starting at 90 dB SPL and decreasing in 5 dB steps. Auditory thresholds were defined as the lowest dB SPL that produced a reproducible ABR trace pattern and were determined manually. All ABR waveform traces were viewed and re-scored by a second operator blinded to genotype. For anesthesia and recovery, see supplemental methods.

### Distortion Product Otoacoustic Emissions

For *Klhdc7b^Regn^* mice, DPOAE measurements were taken on their own or immediately before or after the ABR while the animal was still anesthetized. If the DPOAE was collected on its own, animals were anesthetized as for the ABR recordings.

Two speakers were used to present two primary tones (f1:f2 = 1.2) equally spaced around the three center frequencies measured for ABR (8 kHz, 16 kHz, and 32 kHz). The speakers were used in a closed-field configuration, with tubes connecting the speaker to a sensitive microphone (DPM1, Tucker-Davis Technologies, FL) which was placed inside the ear canal and angled toward the eardrum. Stimuli were calibrated in the ear to be within 2 dB of the target level to account for the effects of the setup. Stimuli were presented with 100 averages. The distortion product was detected at the expected cubic distortion frequency (2f1-f2) and called as positive if the signal was above the noise floor. The noise floor was calculated as the average of five points in the Fourier transform above and below the cubic distortion tone frequency.

For the *Klhdc7b^IMPC^* mice, DPOAE tests were performed at MRC Harwell using frequency-specific tone-burst stimuli at 2 kHz intervals, between 6 and 32 kHz, using the TDT RZ6 System 3 hardware and BioSig RZ (version 5.7.1) software (Tucker Davis Technology). An ER10B+ low noise probe microphone (Etymotic Research) was used to measure the DPOAE near the tympanic membrane. Tone stimuli were presented via separate MF1 (Tucker Davis Technology) speakers, with f_1_ and f_2_ at a ratio of f_2_/f_1_= 1.2 (L1 = 65 dB SPL, L2 = 55 dB SPL), centered around the probing frequencies. Animals were anesthetized as for the ABR recordings. In-ear calibration was performed before each test. The f_1_ and f_2_ tones were presented continuously and a fast-Fourier transform was performed on the averaged response of 356 epochs (each approximately 21 ms). The level of the 2f_1_−f_2_ DPOAE response was recorded and the noise floor calculated by averaging the four frequency bins either side of the 2f_1_−f_2_ frequency.

### Cochlear explant cultures

Cochlear explant cultures were performed with neonatal mice at postnatal days 0-5. The cell culture plates used were 35 mm dishes with a circular punchout and a glass coverslip glued to the bottom (Matsunami, #D35-14-0-U). Collagen bubbles were prepared by mixing 1.7 μl 1 N NaOH, 10 μl 10x PBS, 67 μl rat tail collagen, 21.3 μl H_2_O. 10 μl of this solution was placed on the coverslip and cured at 37°Cin a humidified cell culture incubator with 5% CO_2_ for 30-40 minutes. Collagen bubbles were covered in PBS until use and stored at 4°C. Explant culture media was prepared as follows: 90 ml DMEM/F12 without phenol red, 7 ml fetal bovine serum, 1 ml penicillin G, and 1 ml L-glutamine.

The mice were decapitated and the skull bisected. The cochlea was removed and placed in Leibovitz/L15 media. The bone was opened and the organ of Corti was removed and stria vascularis separated from it. PBS was removed from the culture plate and replaced with cochlear explant media. The organ of Corti was placed on the collagen bubble and most of the media removed to allow the explant to adhere to the collagen. Plates were placed in a humidified cell culture incubator at 37°C with 5% CO_2_. After dissections were finished, 600 μl of culture media was added.

### Gentamycin-Texas Red Assay

Explant cultures were kept in culture and treated 2-3 days later with gentamycin-Texas Red (GtTR, 5 μg/ml) or Texas red (TR, 12.5 μg/ml) diluted in cell culture media. Existing media on the explant was removed, and 600 μL of GTTR or TR containing media was added and cultures were placed in the incubator for 20 minutes. Cultures were rinsed with media, PBS, and then fixed for 15 minutes in 4% PFA, rinsed 3x with PBS and stored at 4°C for several days, after which they underwent immunostaining with the same protocol that was used for whole mount cochlea.

### Statistics

Where applicable, student’s T-test, one or two-way ANOVA as indicated was used to analyze data for significance. All datasets are tested for normality and if failed, non-parametric tests used: Mann-Whitney and Kruskal-Wallis tests. Repeated measures or mixed models were used if data points were missing. If a significant main effect of any variable was observed, Tukey post-hoc tests were performed. Statistics were calculated using GraphPad Prism. * denotes p < 0.05, ** p < 0.01, *** p < 0.001, **** p < 0.0001.

### Variant Effect Predictions and Protein Modelling

Ensembl Variant Effect Predictor (29) was used to generate protein predictions for the effect of the rs36062310 missense variant on KLHDC7B. Alphafold (https://alphafold.ebi.ac.uk/) results for both human isoforms are available with their unique Uniprot IDs (Long variant: A0A3B3ISF6; Short variant: Q96G42). The *.PDB files were downloaded for both the human KLHDC7B isoforms, and these files were utilized for comparative *in-silico* protein structure modelling using [RCSB Protein Data Bank (RCSB PDB)](https://www.rcsb.org/pages/about-us/index) (30). and Missense3D (31).

## Results

### Klhdc7b is expressed in the inner ear and other mouse tissues

To study the role of KLHDC7B in the auditory system we first investigated its expression in the mouse inner ear and several other tissues. Like the human *KLHDC7B* gene, the mouse *Klhdc7b* gene encodes two putative isoforms that differ only in which methionine is annotated as the predicted start codon. It is possible to specifically and individually detect the long isoform transcript [Ensembl transcript [ENST00000648057.3](https://www.ensembl.org/Homo_sapiens/Transcript/Summary?db=core;g=ENSG00000130487;r=22:50545899-50551023;t=ENST00000648057)], but not the short isoform transcript [Ensembl transcript [ENST00000395676.4](https://www.ensembl.org/Homo_sapiens/Transcript/Summary?db=core;g=ENSG00000130487;r=22:50545899-50551023;t=ENST00000395676)] since its sequence is common to both transcripts. We therefore designed two sets of qPCR primers and probes, one hybridizing to the unique long portion of the transcript only (L), and the other probe hybridizing within the region common to both long and short isoforms (L+S) (Fig. 1A top). qPCR data shown in Figure 1 are normalized as ΔCT relative to expression of the housekeeping gene *Drosha*, then represented as Relative Quantification to adult kidney (left) or P1 kidney (right) due to its low expression level in kidney. *Klhdc7b* is expressed at relatively high levels in adult mouse cochlea compared with other tissues (Fig. 1A, left). To assess the expression level of *Klhdc7b* in mouse cochlea over time, we collected cochleae at postnatal days 1, 7, in adult, and aged animals, and confirmed that expression is relatively high throughout life in cochlea, with highest expression in adult cochlea (Fig. 1A, right).

### Klhdc7b is expressed in hair cells

After determining that *Klhdc7b* is expressed in the cochlea via qPCR, we aimed to pinpoint which specific cell types within the inner ear express *Klhdc7b* using *in situ* hybridization and immunofluorescent staining. As in the qPCR probe design (Fig. 1A), two RNAscope probes were designed to detect the common portions of both transcripts (L+S) or a portion unique to the long (L) *Klhdc7b* transcript. We performed RNAscope in conjunction with immunostaining for Myosin VIIa (MYO7A), a hair cell marker (Fig. 1B). Both probes detected transcripts exclusively within hair cells (Fig. 1B), the sensory cells of the cochlea and vestibular system; inner, outer, and vestibular hair cells (IHC, OHC, VHC in Fig. 1B) and were not observed in other cell types. Furthermore, immunostaining with a custom-generated antibody to KLHDC7B is also observed to be exclusively to inner and outer hair cells within the cochlea (Fig. 1C). Staining within hair cells localizes to the plasma membrane, as determined by a co-stain with prestin, a protein localized to the outer hair cell membrane (Fig. 1D).

### Generation of Klhdc7b knockout mice

To investigate the functional role of *Klhdc7b* in the cochlea and to determine whether KLHDC7B is required for hearing, two transgenic mouse lines were generated, *Klhdc7b^RegnΔ/Δ^* and *Klhdc7b^IMPC−/−^,* designed to knock out both the putative long and short isoforms of the gene on different genetic backgrounds. The predominant strain used in transgenic mouse models is the C57BL/6N strain which contains a strain-specific allele (*Cdh23^ahl^*) that confers a progressive age-related hearing loss on these mice (21). *Klhdc7b^IMPC−/−^*mice were generated on the C57BL/6N background (containing the age-related hearing loss mutation in *Cdh23*) as part of the International Mouse Phenotyping Consortium (24) by CRISPR-induced mutation (Fig. 2A, right). This created a 1258 nucleotide deletion within the single exon introducing a premature stop codon and a null allele (32). *Klhdc7b^RegnΔ/Δ^* were generated on the B6.CAST-*Cdh23^753A>G^* background with a 3,787 bp deletion encompassing both the putative long and short isoforms of the gene (Fig. 2A, left).

**Figure 2.**
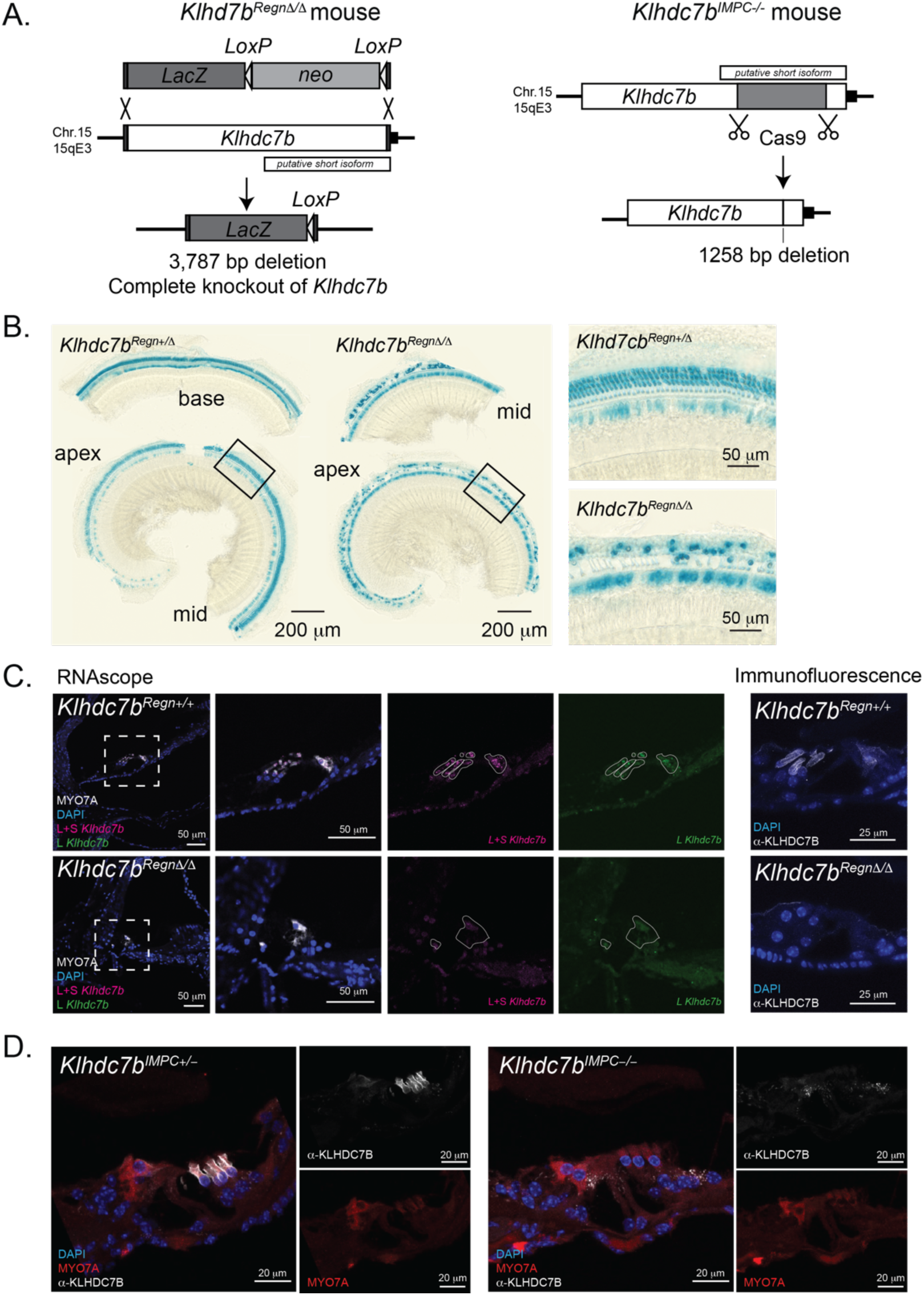
Mutant mouse model design and validation. **A.** Left, schematic showing knockout design for *Klhdc7b^RegnΔ/Δ^* mice. The whole *Klhdc7b* gene was excised from the mouse genome and replaced with a LacZ expression cassette for visualization. Right, knockout design for *Klhdc7b^IMPC−/−^*mice. CRISPR/Cas9 was used to create a 1258 bp deletion creating a truncating premature stop codon and a null allele. The location of the putative short isoform is shown with a smaller box above or below the *Klhdc7b* long isoform in both schematics. **B.** LacZ expression in wholemount preparations of the organ of Corti of heterozygous (*Klhdc7b^Regn+/Δ^*) and knockout (*Klhdc7b^RegnΔ/Δ^*) mice (10-week-old males). **C.** Left, confocal images showing RNAscope *Klhdc7b* probes for the L and L+S transcripts and hair cell marker MYO7A immunofluorescence of *Klhdc7b^Regn+/+^*(top) and *Klhdc7b^RegnΔ/Δ^* mice (bottom), DAPI stains nuclei. Outlines of MYO7A staining are shown overlaid in white on individual probe channels to delineate hair cells. Right, KLHDC7B immunofluorescence of *Klhdc7b^Regn+/+^*(top) and *Klhdc7b^RegnΔ/Δ^* mice (bottom). **D.** Confocal images showing cochleae from *Klhdc7b^IMPC+/−^*(left) and *Klhdc7b^IMPC−/−^* (right) mice at postnatal day 19. Sections have been stained for KLHDC7B (white), MYO7A (red) and DAPI (blue).

Mice for both lines were born at expected Mendelian ratios and appeared phenotypically normal. Since a *LacZ* reporter cassette was inserted within the coding region of *Klhdc7b* in the *Klhdc7b^RegnΔ/Δ^* line, we identified *Klhdc7b* expressing cells using Beta galactosidase (Fig. 2B). At 10 weeks of age, LacZ can be observed in the single row of inner hair cells (IHC) and three rows of outer hair cells (OHC) in cochlear epithelia prepared from *Klhdc7b^RegnΔ/Δ^* and *Klhdc7b^Regn+/Δ^* mice (Fig. 2B). No other cells in the inner ear, including vestibular hair cells, showed LacZ staining (not shown) and no other tissues, including brain and liver, clearly expressed LacZ, which was somewhat surprising since we detected *Klhdc7b* expression using RNA-based techniques (Fig. 1A). Importantly, using both the custom anti-KLHDC7B antibody and RNAScope probes no signal was detected in hair cells of *Klhdc7b^RegnΔ/Δ^* mice (Fig. 2C). *Klhdc7b^IMPC−/−^* mice also showed no presence of anti-KLHDC7B immunostaining in hair cells whereas in *Klhdc7b^IMPC+/−^*mice immunostaining is localized to the membrane of inner and outer hair cells (Fig. 2D) similar to that observed in wildtype mice. (Fig. 1 C and D).

To examine a role for *Klhdc7b* in the central nervous system, we performed a suite of behavioral tests to determine whether there were any changes in exploratory behavior (open field test), spatial working memory (Y-Maze), or motor function and balance (pole test, rotarod). We confirmed that both sexes of *Klhdc7b^Regn+/+^* and *Klhdc7b^RegnΔ/Δ^* mice performed indistinguishably on all four tests (Suppl. Fig. 1). Other measures of phenotypic differences, including weight, bone volume and fat volume, spleen immune profiling, blood immune cell profiling, and serum chemistry were all normal. *Klhdc7b^IMPC−/−^* mice underwent wide-ranging phenotype and behavioral testing as part of the IMPC program, the results of which are available at https://www.mousephenotype.org/data/genes/MGI:3648212. At the start of this study hearing associated abnormalities were the only significant phenotypes recorded in *Klhdc7b^IMPC−/−^* mice. More recently, data released by the IMPC suggest evidence of new additional phenotypes. These include increased eosinophil cell number and decreased lymphocyte cell number in female *Klhdc7b^IMPC−/−^* mice in late adulthood.

### Both Klhdc7b knockout mouse models show early onset progressive hearing loss

Given its specific expression in cochlear hair cells, we studied the role of KLHDC7B in the inner ear, by measuring Auditory Brainstem Responses (ABR) in wildtype, heterozygous and homozygous mutant mice for both *Klhdc7b^RegnΔ/Δ^* (Fig. 3A) and *Klhdc7b^IMPC−/−^* (Fig. 3B) knockout models at several ages. ABRs measure the electrical activity of the auditory pathway in response to a sound stimulus as the electrical signal travels from the cochlear nerve up through the brainstem toward the auditory cortex. Hearing function in mice begins at two weeks of age. At postnatal day 17 (P17, 2-3 weeks), just after the onset of hearing, *Klhdc7b^RegnΔ/Δ^* mice exhibit significantly elevated ABR thresholds compared to their littermates (Fig. 3A, 2-3 weeks), indicating a severe hearing loss (Fig 3A, left). Additionally, the *Klhdc7b^RegnΔ/Δ^* mice have a significantly diminished Wave 1 amplitude compared to their littermates (Suppl. Fig. 2), which may be reflecting differences in thresholds. ABR thresholds become progressively elevated as the *Klhdc7b^RegnΔ/Δ^* mice age, such that most have no response at any of the three tested frequencies by 11-15 weeks of age (Fig. 3A, right).

**Figure 3.**
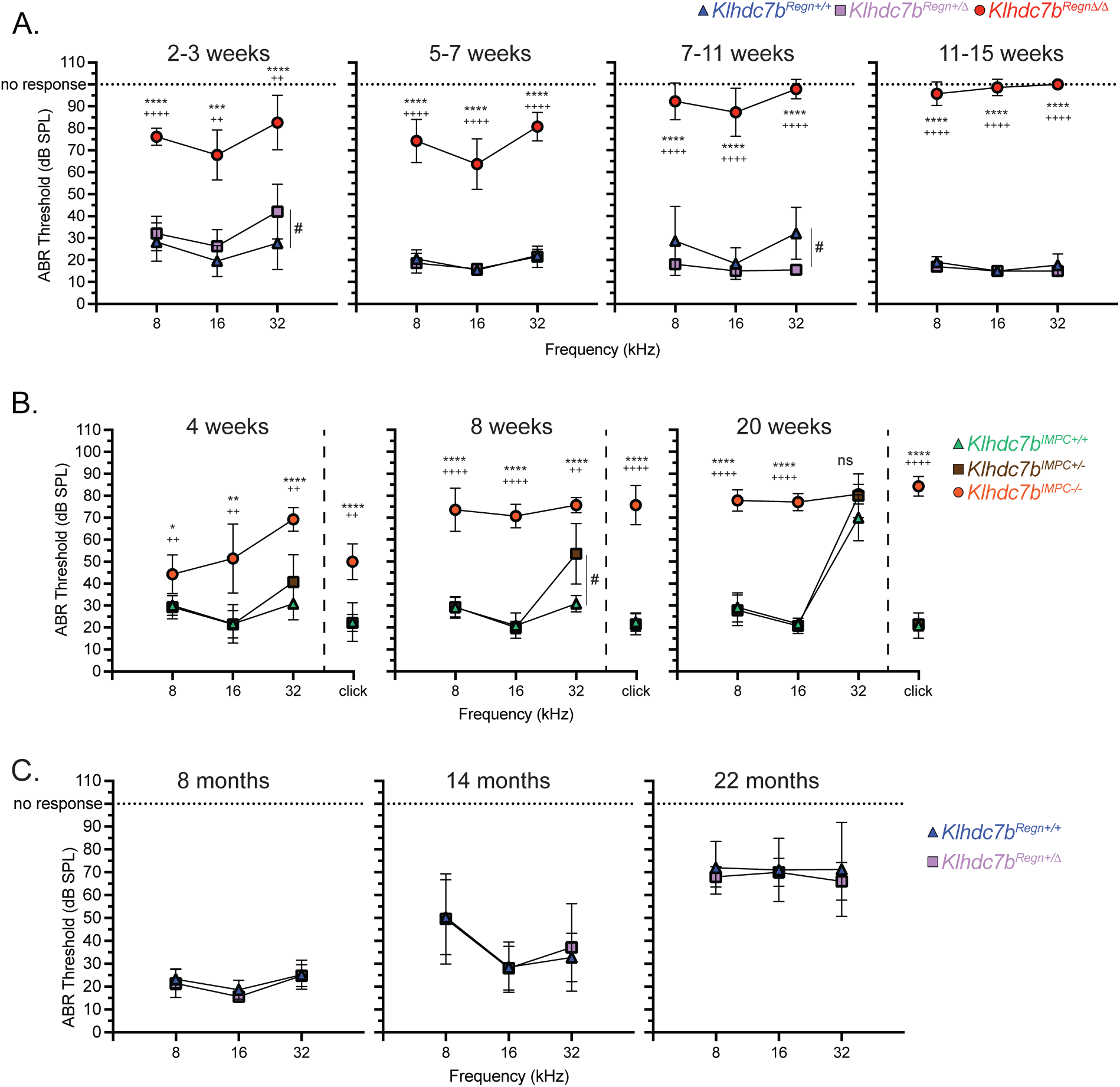
*Klhdc7b-*deficient mice exhibit early onset progressive hearing loss. **A.** ABRs were recorded in separate cohorts of *Klhdc7b^Regn+/+^*, *Klhdc7b^Regn+/Δ^* and *Klhdc7b^RegnΔ/Δ^* mice at four time points. 100 dB denotes no response. **B.** ABRs were recorded longitudinally in a cohort of *Klhdc7b^IMPC+/+^, Klhdc7b^IMPC+/−^* and *Klhdc7b^IMPC−/−^*mice at three time points. **C.** ABRs were recorded longitudinally in a cohort of *Klhdc7b^Regn+/Δ^* and *Klhdc7b^RegnΔ/Δ^* mice at three older time points. For A-C, mean ABR (±SD) thresholds for click and pure-tone response are shown. Analysis was performed via two-way ANOVA, using Tukey’s test for post-hoc comparisons. For A-C * p<0.05, ** p<0.01, *** p<0.001, **** p<0.0001 between knockout (*Klhdc7b^RegnΔ/Δ^, Klhdc7b^IMPC−/−^)* compared with wild type (*Klhdc7b^Regn+/+^*, *IMPC^+/+^*) mice. + p<0.05, ++ p<0.01, +++ p<0.001, ++++ p<0.0001 between knockout and heterozygous (*Klhdc7b^Regn+/Δ^, Klhdc7b^IMPC+/−^)* mice. # p <0.05 between wild type and heterozygous mice.

Similarly, *Klhdc7b^IMPC−/−^* mice exhibit elevated ABR thresholds at 4 weeks of age, the youngest age tested, showing moderate hearing loss at lower frequencies progressing to severe at 32 kHz, the highest frequency tested (Fig. 3B). In *Klhdc7b^IMPC−/−^* mice, hearing loss progresses with age reaching a ~80dB SPL threshold by 20 weeks of age across all frequencies tested. At this age the effect of the *Cdh23^ahl^* allele in the C57BL/6N strain is becoming evident, with significantly elevated high-frequency (32 kHz) ABR thresholds also observed in heterozygote and wildtype mice (Fig. 3B).

For both *Klhdc7b* knockout models there was no overt hearing loss phenotype in heterozygote mice although small significant elevation of thresholds compared to wildtype mice were found at 32 kHz at some time points (Fig. 3A left, 3B middle). To test whether there might be an age-related phenotype in heterozygous mice, *Klhdc7b^Regn+/Δ^* mice were aged along with *Klhdc7b^Regn+/+^* controls. ABR thresholds (Fig. 3C) and Wave 1 amplitudes (Suppl. Fig. 3) were indistinguishable from *Klhdc7b^Regn+/+^* mice as late as 22 months of age. Interestingly, heterozygous *IMPC^+/−^*mice develop high-frequency hearing loss at 32 kHz earlier than their wildtype counterparts, having higher thresholds at 8 weeks of age (Fig 3B, middle).

To further characterize the hearing phenotype of these mice, we recorded DPOAE to assess the *in vivo* function of outer hair cells and found that there was remaining OHC function in *Klhdc7b^RegnΔ/Δ^* mice at 2-3 weeks, with very little remaining by 12 weeks when the ABR signal has disappeared (Fig. 4A, right). To better understand the time course of the loss of OHC function, a cohort of *Klhdc7b^RegnΔ/Δ^* and *Klhdc7b^Regn+/Δ^* littermates were longitudinally recorded every week from 2-3 weeks to 12 weeks of age. There is some remaining OHC function at 2-3 weeks of age, with DPOAE thresholds becoming significantly elevated in *Klhdc7b^RegnΔ/Δ^* mice compared with *Klhdc7b^Regn+/Δ^* from 5 weeks of age (Fig. 4B). When tested at 20 weeks of age, *Klhdc7b^IMPC−/−^* mice exhibit a similar loss of OHC function, as observed in *Klhdc7b^RegnΔ/Δ^* mice, whereas their wildtype and heterozygote littermates continue to show good low frequency DPOAE responses (Fig 4C).

**Figure 4.**
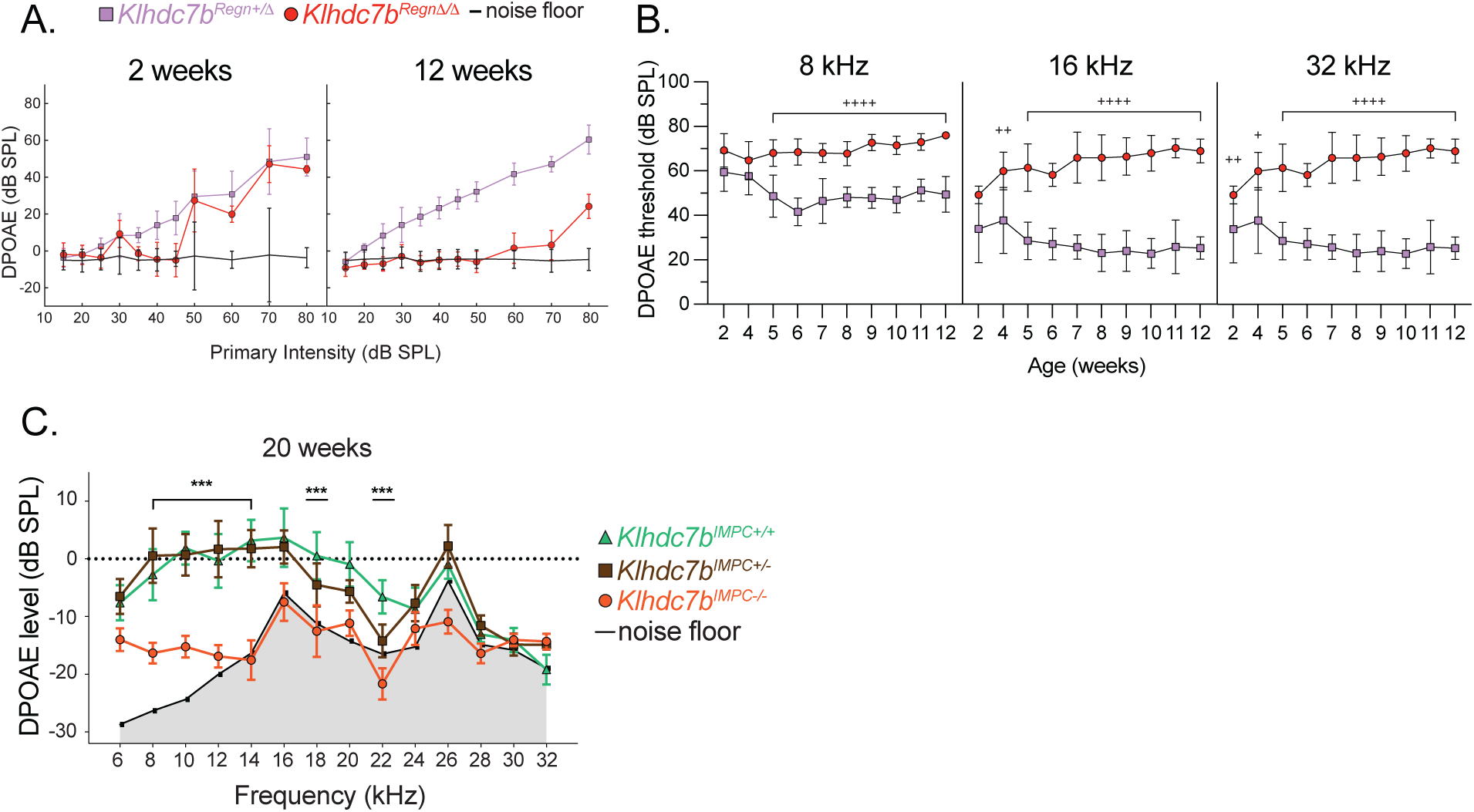
DPOAE in *Klhdc7b-*deficient mice indicate outer hair cell dysfunction. **A,B.** DPOAE longitudinally recorded in a cohort of *Klhdc7b^Regn+/+^*, *Klhdc7b^Regn+/Δ^* and *Klhdc7b^RegnΔ/Δ^* mice. **A**. Input-output curves of DPOAE signal plotted against primary intensity of presented stimulus at 2 and 12 weeks for 16KHz. **B.** DPOAE thresholds over time plotted at three frequencies. **C.** DPOAEs recorded in *Klhdc7b^IMPC+/+^, Klhdc7b^IMPC+/−^,* and *Klhdc7b^IMPC−/−^*mice plotted against frequency. The stimulus was presented in 2 kHz intervals at 65/55 dB SPL for the f1 and f2 tones, respectively. For A-B, Analysis was performed via two-way ANOVA, using Tukey’s test for post-hoc comparisons. For C, Analysis was performed via two-way ANOVA followed by unpaired t-tests for each frequency. For A-C, * p<0.05, ** p<0.01, *** p<0.001, **** p<0.0001 between knockout (*Klhdc7b^RegnΔ/Δ^, Klhdc7b^IMPC−/−^)* compared with wild type (*Klhdc7b^Regn+/+^*, *Klhdc7b^IMPC+/+^)* mice. + p<0.05, ++ p<0.01, +++ p<0.001, ++++ p<0.0001 between knockout and heterozygous (*Klhdc7b^Regn+/Δ^, Klhdc7b^IMPC+/−^*) mice.

### Outer hair cells in both Klhdc7b knockout mouse models develop normally, but begin to die around hearing onset

The hearing phenotype of *Klhdc7b^RegnΔ/Δ^* and *Klhdc7b^IMPC−/−^* mice led us to examine the anatomy of the organ of Corti in these mice. Examples of middle-apical turn of the organ of Corti of *Klhdc7b^RegnΔ/Δ^* and *Klhdc7b^Regn+/+^* at different ages are shown in Figure 5A. At P6, the gross anatomy of the organ of Corti of *Klhdc7b^RegnΔ/Δ^* mice appear normal, immunostaining with hair cell marker MYO7A shows that the 3 rows of outer hair cells and single row of inner hair cells are intact and stereocilia are present, stained with F-actin, (shown in examples from the middle turn in Figure 5A). However, by p11 a few MYO7A+ outer hair cells appear to be missing (Fig 5A, white arrows). By 3 weeks, many outer hair cells are missing and there are visible supporting cell scars (Fig 5A, white arrowheads), which are known to form when hair cells die (33). Circular stained areas of MYO7A (Fig. 5A, asterisks) are present beyond the normal border of the hair cell location, which may suggest engulfment of dead hair cells by supporting cells (34, 35). By 8 weeks of age, almost no outer hair cells are present and evident scarring is visible as a lattice-like pattern of F-actin staining (Fig. 5A, arrowheads). This outer hair cell death phenotype is consistent with the increased thresholds observed in the ABR and DPOAE recordings.

**Figure 5.**
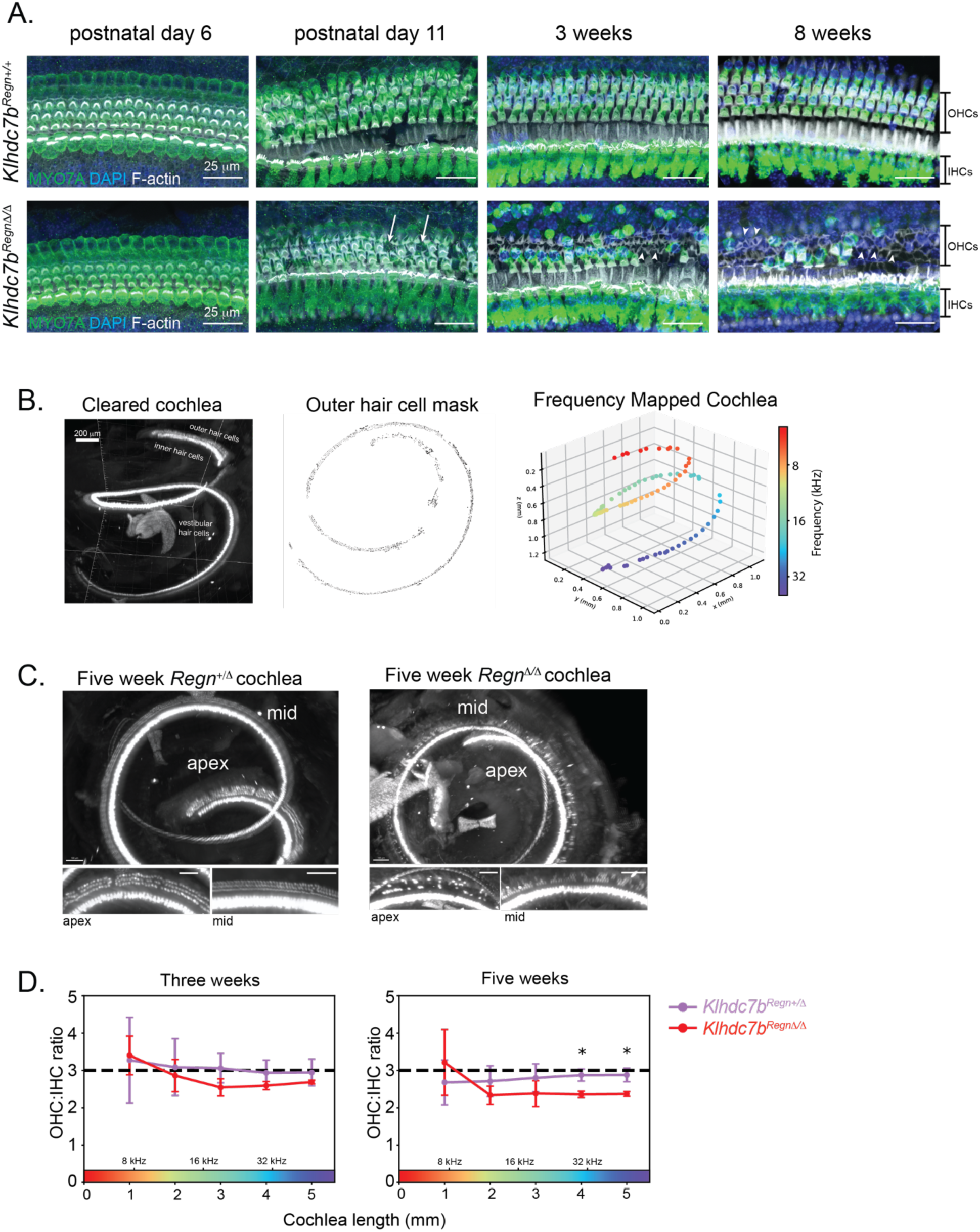
Outer hair cell death in *Klhdc7b^RegnΔ/Δ^* mice. **A.** Immunostaining of whole mounted cochlea of *Klhdc7b^Regn+/+^*, and *Klhdc7b^RegnΔ/Δ^* mice at postnatal day (p) 6, p11, 3 weeks and 8 weeks of age. MYO7A (green) specifically labels hair cells, DAPI (blue) stains nuclei, F-actin (white) labels stereocilia. Arrows denote missing OHCs. Arrowheads denote examples of supporting cell scars where OHCs have died. Asterisks denote rounded MYO7A staining of cells outside the normal hair cell location. **B.** Cochleae were stained for MYO7A and a nuclear stain, then cleared and imaged using light-sheet microscopy. For one example cochlea, Left, MYO7A staining in 3D. Middle, automatically generated outer hair cell mask. Right, frequency mapping along the tonotopic axis from apex to base. **C.** Representative example images of *Klhdc7b^Regn+/Δ^* (left) and *Klhdc7b^RegnΔ/Δ^* (right) cochleae at five weeks visualized from the top down. Below each image of intact cochlea are example images of apex (left) and middle (right) turns. **D.** Outer to inner hair cell ratios at three weeks and five weeks from *Klhdc7b^RegnΔ/+^* and *Klhdc7b^RegnΔ/Δ^* cochleae. n=3 mice per genotype, two-tailed Welch’s t-test, *p<0.05

Additionally, we quantified hair cells in intact cochlea using light-sheet microscopy to better visualize all regions of the cochlea. This method has been previously published in gerbil (27). We adapted the protocol for mouse and utilized the anti-MYO7A antibody and light-sheet microscopy to visualize hair cells (Fig. 5B). Inner and outer hair cell masks were automatically annotated and projected onto the frequency map of the mouse cochlea. A clear reduction in outer hair cells was apparent at 5 weeks (Fig. 5C). At three weeks of age, the ratio of outer to inner hair cells along the length of the cochlea is slightly smaller in *Klhdc7b^RegnΔ/Δ^* than in *Klhdc7b^Regn+/Δ^* mice, and the difference is significant toward the base of the cochlea at five weeks of age (Fig. 5D).

The *Klhdc7b^IMPC−/−^* mouse model also showed a similar progressive outer hair loss (Fig. 6 and Suppl. Fig. 4). At early postnatal time points (P2, Fig. 6A), staining for MYO7A reveals both the 3 rows of outer hair cells and single row of inner hair cells appeared normal with no significant difference in hair cell number between all genotypes. At 3 weeks, there was significant outer hair cell loss at the base of the cochlea (Fig. 6B), but not in the mid or apical turns. By 8 weeks old this had progressed to significant outer hair cell loss at all portions of the cochlea, with no inner hair cell loss at this age (Fig. 6C and D). The timing of outer hair cell death corresponded to the progressive hearing loss shown in Figure 3 for both models. Suppl. Figure 4 shows higher resolution images with immunostaining of supporting cell marker Sox2 and F-actin labelled stereocilia in which morphology of the non-sensory epithelium and stereocilia appear similar in *Klhdc7b^IMPC−/−^, Klhdc7b^IMPC+/−^* and *Klhdc7b^IMPC+/+^*at early ages.

**Figure 6.**
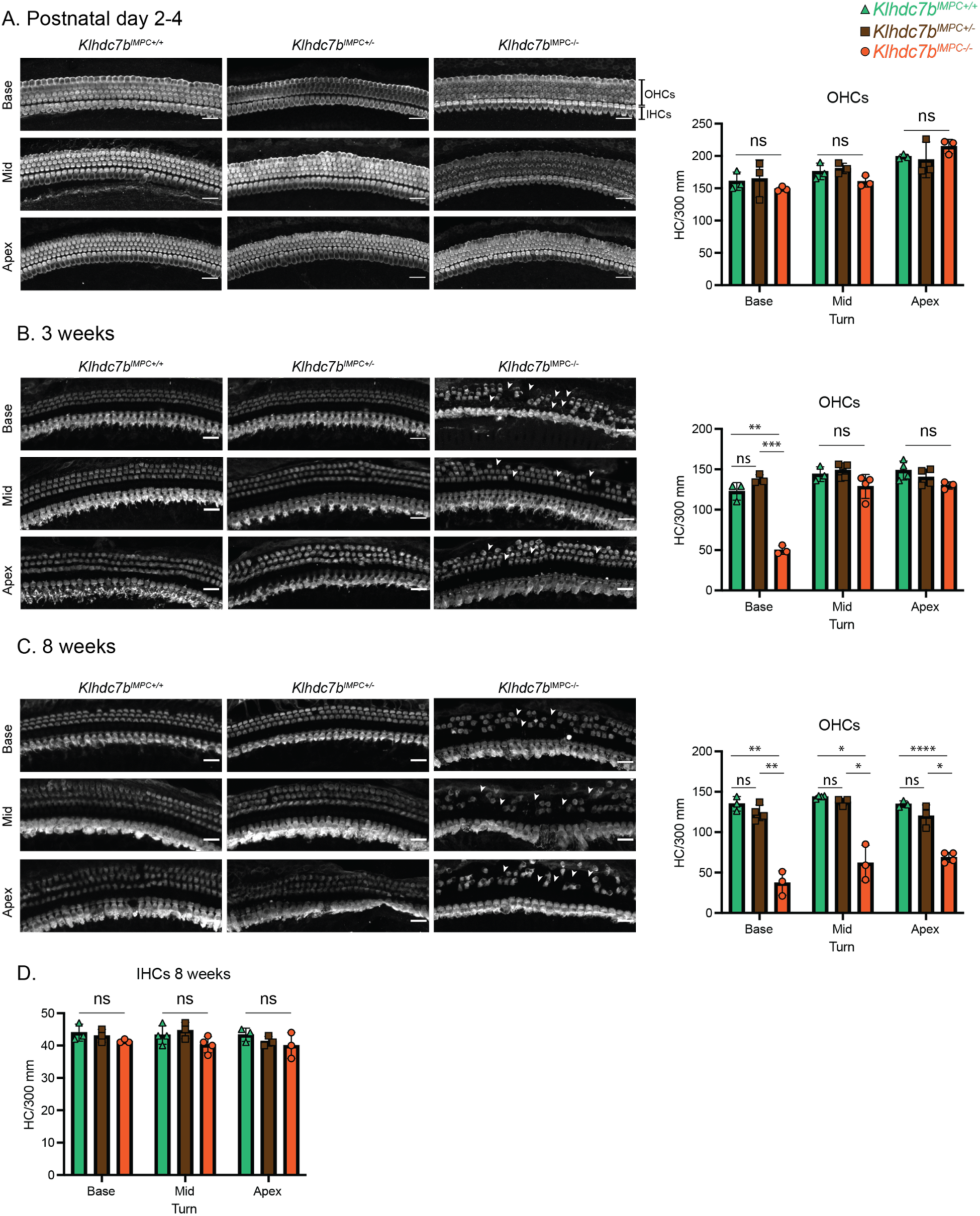
Progressive outer hair cell loss in *Klhdc7b^IMPC−/−^* mice is evident from 3 weeks of age. **A-C**, Immunofluorescence in wholemount cochlea preparations using MYO7A to label hair cells in *Klhdc7b^IMPC+/+^, Klhdc7b^IMPC+/−^* and *Klhdc7b^IMPC−/−^* mice. Panels show (x20) representative confocal microscopy images from all three genotypes and charts to the right showing OHC quantification at each cochlea turn, base, mid and apical. Scale bar = 5 µm. Data is shown for 3 ages: (*A*) postnatal day 2-4; (*B*) 3 weeks and at 8 weeks of age (*C*). Arrowheads show examples of missing outer hair cells. **D.** Chart shows quantification of inner hair cells (IHCs) at 8 weeks of age. Statistical analysis was performed via one-way ANOVA, using Tukey’s test for post-hoc comparisons, *ns*, not significant, * p<0.05, ** p<0.01, *** p<0.001, **** p<0.0001.

### Outer hair cells appear to develop normal stereocilia and functional mechanotransduction channels

Deficits in stereocilia formation or morphological abnormalities can lead to deafness and hair cell death (36–40). To determine whether the hair cell degeneration was driven by stereocilia abnormalities, we performed SEM and TEM in organ of Corti from our mutant mouse models. SEM in *Klhdc7b^Regn+/Δ^* and *Klhdc7b^RegnΔ/Δ^* mice at postnatal days 10 (P10) and 3 weeks were undertaken to visualize the organ of Corti at time points before and during outer hair cell degeneration. Sample images from the middle turn of the organ of Corti are pictured in Fig. 7A. Both *Klhdc7b^Regn+/Δ^* mice and *Klhdc7b^RegnΔ/Δ^* mice have stereocilia that appear grossly normal at P10 (Fig. 7A, left). By 3 weeks, many outer hair cells are absent, however, the stereocilia bundles on the remaining outer hair cells are intact with apparently normal morphology (Fig. 7A, right). Similarly, SEM data demonstrated the stereocilia bundle of most OHC appears normal in *Klhdc7b^IMPC−/−^* mice at 3 weeks of age (Fig. 7B, left). By 8 weeks and 25 weeks of age some stereocilia pathology is visible in surviving outer hair cells of *Klhdc7b^IMPC−/−^* mice, potentially due to hair cells being in the process of dying (Fig. 7B, middle and right). Transmission electron microscope (TEM) images also suggest IHC and OHC stereocilia appear normal in mice at 3 weeks of age (Fig. 7C). These data suggest that KLHDC7B is not critical for the development and integrity of the stereocilia bundle.

**Figure 7.**
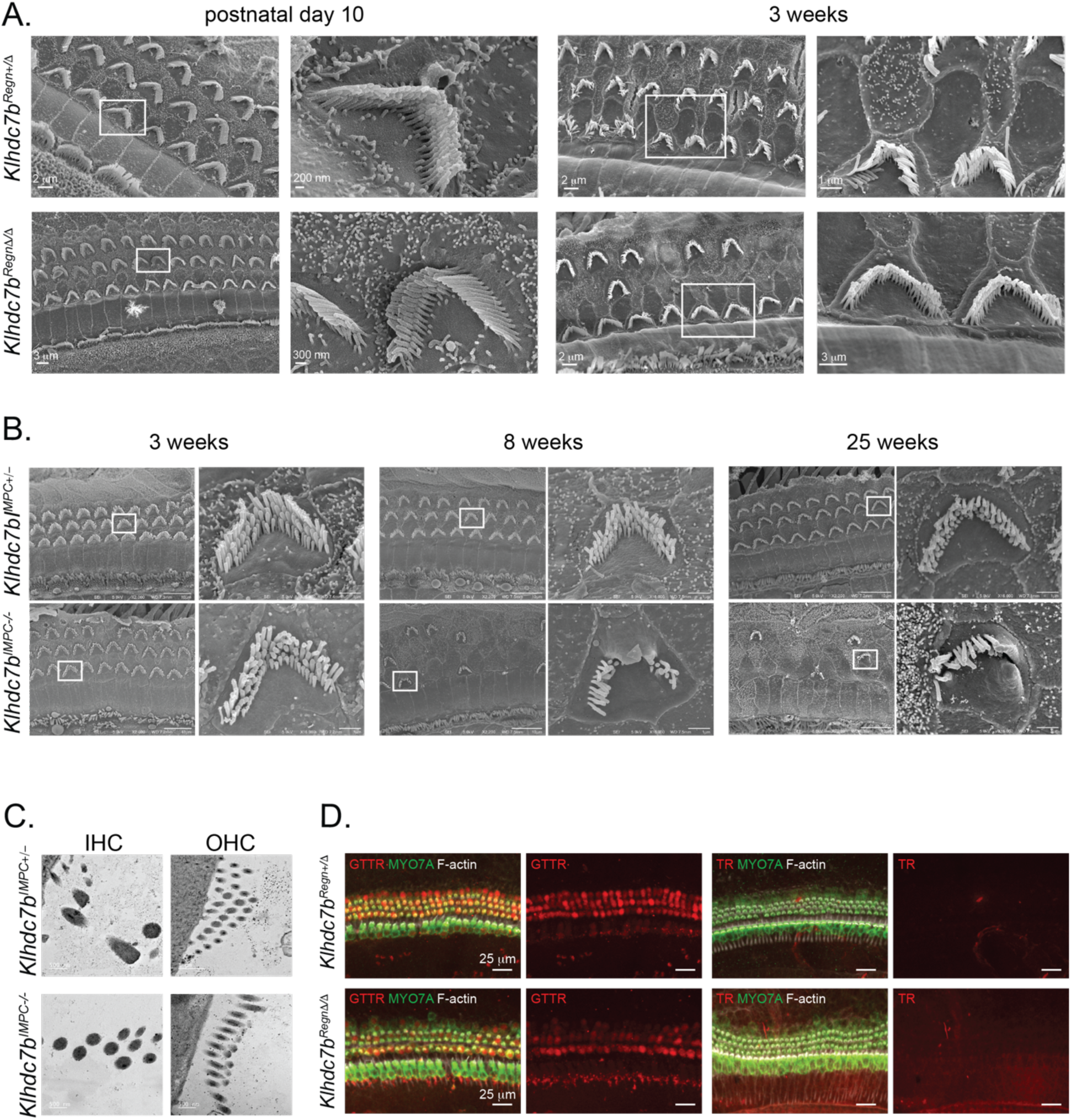
Stereocilia and mechanotransduction channel function appear normal in *Klhdc7b-*deficient mice. **A.** SEM of *Klhdc7b^RegnΔ/+^* and *Klhdc7b^RegnΔ/Δ^* cochlea at P10 and 3 weeks. Stereocilia appear grossly normal, even after significant hair cell loss at 3 weeks. **B.** SEM of *Klhdc7b^IMPC+/−^* and *Klhdc7b^IMPC−/−^* cochlea at 3 weeks (left), 8 weeks (middle), and 25 weeks (right) of age show normal development of stereocilia. Scale bar in left images at each time point, 10 μm, in cutout images 1 μm. **C.** TEM of stereocilia of *Klhdc7b^IMPC+/−^,* and *Klhdc7b^IMPC−/−^* cochlea at 3 weeks suggest IHCs and OHCs have normal stereocilia. **D.** Cochlear explant cultures of *Klhdc7b^RegnΔ/+^* and *Klhdc7b^RegnΔ/Δ^* mice treated with Gentamycin-Texas Red (GtTR, left) or Texas Red (TR, right) alone, then immunostained for MYO7A labelling hair cells and stained for F-actin (stereocilia). Entry of GtTR into hair cells confirms mechano-transduction in both *Klhdc7b^RegnΔ/+^*and *Klhdc7b^RegnΔ/Δ^* mice.

Mechanotransduction (MET) defects can also cause hair cell death (41), so we sought to assess mechanotransduction (MET) complex function using a gentamycin-Texas red assay (GtTR). Gentamycin, like other aminoglycosides, is known to readily enter hair cells via the MET complex and this property is maintained when conjugated to a fluorescent dye, Texas red (42–45). In the absence of functional MET complexes GtTR will not enter hair cells thus allowing us to assess hair cell function before the onset of hearing in mice. To evaluate this, we cultured neonatal organ of Corti from *Klhdc7b^RegnΔ/Δ^* and *Klhdc7b^Regn+/+^* mice at P4-P5 for 3 days before incubating with GtTR for 20 minutes. GtTR labeling was observed in hair cells for both the *Klhdc7b^RegnΔ/Δ^* and *Klhdc7b^Regn+/+^* cultures, indicating that MET complexes are assembled and functional (Fig. 7D).

## Discussion

The auditory system is highly complex, encompassing multiple tissues and many specialized cell types. This is evidenced from the high heterogeneity of functions and cell-specific expression of genes associated with monogenic hearing loss, with several hundred loci found to date (46, accessed 20 Sept 2024). Although good progress has been made in understanding monogenic hearing loss, the genes and hence pathogenic mechanisms that underlie the highly prevalent ARHL have been much more difficult to identify.

Early GWAS with well-characterized ARHL cohorts largely did not identify genome-wide significant associations (5). Recent GWAS with much larger cohorts have been more successful, identifying over 50 associations, but these have largely relied on self-report hearing data or medical records rather than more reliable audiometric measures to define the phenotype. Therefore, some doubt remained about whether these associations represent true ARHL susceptibility genes. Our results here demonstrate that *Klhdc7b*, the ortholog of a novel gene strongly associated with hearing loss in several recent human GWAS focused on age-related hearing loss, is necessary for the maintenance of hearing in mice. This is the first example of a novel gene identified in a genetic association study for age-related hearing loss to be validated in mice and provides a strong corroboration of the findings of recent GWAS using self-reported hearing loss data.

We find that *Klhdc7b* is expressed in the cochlea of mice, exclusively in hair cells (Fig. 1). We demonstrate that loss of *Klhdc7b* in mice causes a severe, progressive hearing loss in two independent *Klhdc7b* knockout mouse models (Fig. 3). This loss occurs from hearing onset, with a greater hearing deficit at the higher frequencies than lower frequencies that is more notable in the *Klhdc7b^IMPC−/−^* model. In both models, the hearing loss progresses to a profound or complete loss at all frequences tested within a few weeks. The cochlea is organized tonotopically, with high frequencies detected at the basal end of the cochlea, whereas low frequencies are detected towards the apex (47). Histological assessments in whole mount preparations of cochlea from the *Klhdc7b* knockout mouse models reveal a progressive loss of outer hair cells proceeding from base to apex, corresponding to the hearing deficit. Inner hair cells remain intact as of 8 weeks of age (Fig. 5A, 6D).

Our data suggest that *Klhdc7b* is not necessary for the development and functional maturation of hair cells, including the arrangement of their stereocilia or the assembly of functional mechanotransduction complexes (Fig. 7). The observation that the *Klhdc7b^IMPC−/−^* model still retains some hearing particularly at lower frequencies at 1 month of age suggests that the hair cells that develop are functional. The remaining DPOAE at early ages in the *Klhdc7b^RegnΔ/Δ^* model also suggests the hair cells that develop are functional (Fig. 4). Taken together, these results suggest that *Klhdc7b* is required for maintenance and survival of outer hair cells.

The length and stiffness of outer hair cells varies along the length of the cochlea, allowing outer hair cells to be frequency-selective (48). Outer hair cells are known to be more sensitive than inner hair cells, dying earlier than inner hair cells in response to insults such as drug-induced toxicity and noise exposure (49–51). This generally occurs in a base-to-apex pattern, perhaps due to differences in the properties of hair cells or differential exposure to stress along the length of the cochlea. This base-to-apex pattern is observed in early stages of ARHL, later progressing to inner hair cell loss (52–54). Both *Klhdc7b* knockout models exhibit a base-to-apex pattern of outer hair cell loss (Fig. 5, 6). The causes of outer hair cell death are known to be diverse; deficits in electromotility (55), specific calcium buffering proteins (56, 57), autophagy (58), and ER stress regulation (59) can all cause outer hair cell death and hearing loss. Therefore, the requirement of *Klhdc7b* for the survival of outer hair cells does not necessarily provide clues as to its function.

While KLHDC7B does not appear to be required for the survival of inner hair cells and we have found no evidence of inner hair cell dysfunction in our *Klhdc7b* mouse models up to 8 weeks of age, our data do not rule out a functional inner hair cell deficit. However, other mouse models that have specific outer hair cell dysfunction or loss show decreases in ABR thresholds that are consistent with our model (55, 60, 61). According to our data, KLHDC7B is expressed in inner hair cells (Fig. 1), so presumably it has some role in these cells. Perhaps KLHDC7B performs a similar function in inner and outer hair cells, but its function is more critical for outer hair cell survival.

Although we have confirmed the need for KLHDC7B for maintenance of auditory function we have not identified the molecular function of KLHDC7B in the inner ear. There is very little published information on the role and molecular function of KLHDC7B. We show that KLHDC7B appears to be localized to or near the plasma membrane of hair cells, with stronger staining observed in outer hair cells. While other members of the Kelch domain-containing superfamily have diverse cellular locations and functions (18, 19), some do bind to actin, and many are involved in cell structure and organization by acting as scaffolds for protein-protein interactions (18). KLHDC7B also has a predicted transmembrane domain close to the N terminus of the putative long isoform of both the mouse and human protein (amino acids 26-47 in both) according to Phobius, a web-based protein domain prediction tool (62, 63).

The structure near the plasma membrane in outer hair cells is complex. Just inside the plasma membrane of outer hair cells are complex structures called the cortical lattice and subsurface cisternae. The cortical lattice links the subsurface cisternae with the plasma membrane, and it consists of actin molecules cross-linked by spectrin beta V. There are two unknown structural components in this region, the “pillar” linking the cortical lattice with the plasma membrane, and the actin-subsurface cisterna connection that links the actin within the CL to the SSC (64). Further research is needed to determine whether KLHDC7B could be involved in these structures or related to their function.

The two *Klhdc7b* knockout models were generated using independent strategies and on different mouse strain backgrounds (Fig. 2). Despite this, the phenotypes of both knockout lines are remarkably consistent although with some minor differences. Surprisingly, the hearing deficit in the *Klhdc7b^IMPC−/−^* mice on the *Cdh23^ahl^* background is less severe and has a slightly later onset than that found in the *Klhdc7b^RegnΔ/Δ^* mice on the B6.CAST.*Cdh23^753A>G^* background (Fig. 3). This suggests that there is no evidence of an interaction between the *Cdh23^ahl^* and the *Klhdc7b* to produce an earlier onset phenotype as has been found in other hearing gene models (59, 65). It is possible that variations in the conditions of animal housing environments (e.g. noise), and ABR measurements between the two host organizations might account for these minor differences.

In both models, loss of *Klhdc7b* in mice resulted in an early onset and fast progressing hearing loss, whereas data from human GWAS has suggested that variation in *KLHDC7B* is associated with increased risk of ARHL, a much later onset phenotype. This may be accounted for by the different effects of a null allele compared to those of a missense variant detected in the GWAS. However, the requirement of KLHDC7B for survival rather than development of outer hair cells is consistent with a role in ARHL, and in early stages of ARHL progressive outer hair cell loss occurs before inner hair cells (66).

The lack of a phenotype in heterozygous *Klhdc7b^Regn+/Δ^* mice suggests that loss of one copy of the *Klhdc7b* gene is not deleterious to outer hair cell survival in mice. This is of interest since some rare heterozygous loss of function variants in KLHDC7B have been associated with ARHL in human GWAS (17). However, a heterozygous mutation that results in hair cell degeneration over the course of decades in humans exposed to noise and other ototoxic insults may not present within the 2.5-year lifespan of the mouse without the same environmental exposures. It would be interesting to test whether *Klhdc7b^Regn+/Δ^* mice are more susceptible to outer hair cell loss in response to an insult such as noise exposure. *Klhdc7b* expression has been shown to increase in outer hair cells after noise-induced hearing loss (67), which could be a compensatory response to injury.

The variant in *KLHDC7B* which has been consistently and strongly associated with ARHL in recent GWAS is a missense variant which causes a valine to methionine change in both long and short forms of KLHDC7B (15, 17, 68, 69). Interestingly, both Polyphen and Sift predictions suggest that this variant would be deleterious to the long version of the protein but not to the short isoform (Suppl. Table 1). Protein modeling (Suppl. Fig. 5) indicates that the additional alpha helices in the longer form of KLHDC7B may interact and alter the β-propeller structure common to both long and short forms suggesting the two isoforms might have subtly distinct β-propellers. Missense 3D (31) also indicates that the 1154Val>Met change in the long form is structurally damaging (Suppl. Fig. 6). Our expression studies show that the longer form of KLHDC7B is expressed in hair cells in mice. The 1154 valine amino acid is conserved in the mouse gene meaning it should be possible to create a knock-in mouse model of the human Val>Met mutation to investigate the effect on outer hair cell survival.

## Conclusions

Overall, this work supports the use of large-scale GWAS studies to identify novel genetic associations with hearing loss, even when these studies do not have comprehensive auditory phenotyping. Our results translate a finding from a genome-wide association study in humans back into mice, allowing us to fully study the function of the gene in a controlled manner in a model system where the inner ear is much more accessible than in humans. We were able to identify the cellular expression pattern, subcellular location, and characterize the phenotype in two knockout mouse models of *Klhdc7b*. The very similar phenotype in two mouse models generated and analyzed at two different institutions and mouse facilities, possibly with different noise levels, on different backgrounds with respect to a known age-related hearing loss mutation, strengthens the case for a specific hearing loss phenotype caused by the loss of KLHDC7B. The cellular phenotype was characterized by normal development of hair cells with functional mechanotransduction complexes, followed by outer hair cell death correlated with severe and progressive hearing loss. Further work is needed to clarify the molecular function of KLHDC7B.

Age-related hearing loss can not only cause social isolation and a decreased quality of life but is also the largest modifiable risk factor for dementia (70). This emphasizes the importance of studies identifying and characterizing novel risk factors for age-related hearing loss. This study provides a strong rationale to pursue other novel candidates identified by recent ARHL focused GWAS. These studies will also provide a better understanding of the physiology of the inner ear, providing new approaches to help prevent or ameliorate hearing loss in the future.

## List of abbreviations

ARHL: age-related hearing loss
*ahl*: a known age-related hearing loss causing allele in *Cadherin 23* found in C57BL6/N and C57BL6/J mice
ABR: auditory brainstem response
ANOVA: analysis of variance
BAC: bacterial artificial chromosome
bp: base pairs
°C: degrees Celsius
cDNA: complementary deoxyribonucleic acid
CO_2_: carbon dioxide
CRISPR: clustered regularly interspaced short palindromic repeats
DAPI: 4′,6-diamidino-2-phenylindole
dB: deciBels
dB SPL: deciBels sound pressure level
ddH_2_O: double distilled water
DNA: deoxyribonucleic acid
DPOAE: distortion product otoacoustic emission
ECi: Ethyl Cinnamate
EDTA: Ethylenediaminetetraacetic acid
ES: embryonic stem
F0: first generation of mice
Freq: frequency
FWHM: full-width-at-half-maximum
GtTR: gentamycin Texas Red
GWAS: genome-wide association study
IHC: inner hair cell
IMPC: International Mouse Phenotyping Consortium
kHz: kilohertz
KLHDC7B: Kelch domain containing 7B
KO: knockout
LacZ: beta galactosidase
MET: mechanotransduction complex
mRNA: messenger ribonucleic acid
ms: milliseconds
MYO7A: myosin VII A
OHC: outer hair cell
P: postnatal day
PBS: phosphate buffered saline
PBS-T: phosphate buffered saline plus triton X-100
PFA: paraformaldehyde
qPCR: quantitative polymerase chain reaction
RNA: ribonucleic acid
Rpm: rotations per minute
Rt-PCR: reverse transcriptase polymerase chain reaction
SDS: sodium dodecyl sulfate
SEM: scanning electron microscopy
TEM: transmission electron microscopy
VHC: vestibular hair cell
w/v: weight per volume

## Declarations

Ethics approval and consent to participate: Not applicable

Consent for publication: Not applicable

Availability of data and materials: The datasets used and/or analyzed during the current study are available on reasonable request to the authors. The *Klhdc7b^IMPC−/−^*mice are available from the IMPC.

## Competing interests

AMK, RAD, DJ, AC, BJvS, NV, GB, LCB, JC, EF, NZ, ML, SC, KC, DDBM, JM, SDC, JRW, MG, MCD are all employees of Regeneron Pharmaceuticals, Inc. BS, CA, LN, AB, MRB, SJD have no competing interest.

## Funding Sources

Benjamin Silver’s PhD was funded by the Geoffrey Duveen Research Fellowship in Otology and the Royal College of Surgeon of England Research Fellowship. This work was supported by Dunhill Medical Trust Medical (RPGF2002\189 and CER\91 to SJD), RNID (F98 to SJD) and the Medical Research Council (MC_UP_1503/2 and MR/X004597/1 to MRB).

## Authors’ contributions

AMK, BS, MRB, SDC, MCD, and SJD designed and planned experiments. AMK, BS, CA, LN, DJ, AB, AC, GB, LCB, JC, EF, SC, KC, DDBM, JM performed experiments. RAD designed the *Klhdc7b^RegnΔ/Δ^* knockout mouse model. BJvS, NV, JRW, MG advised and analyzed data for 3D imaging experiments. CA, DJ, NZ, and ML analyzed electrophysiological data (ABR and DPOAE). AMK and SJD wrote the manuscript. All authors reviewed and edited the manuscript.

## Acknowledgements

The authors acknowledge Irina Marcovich for helpful comments on the manuscript and the Ear Institute microscopy unit for help with imaging.

