## Supplemental Data for "KLHDC7B, a novel gene associated with age-related hearing loss in humans, is required for the maintenance of hearing in mice"

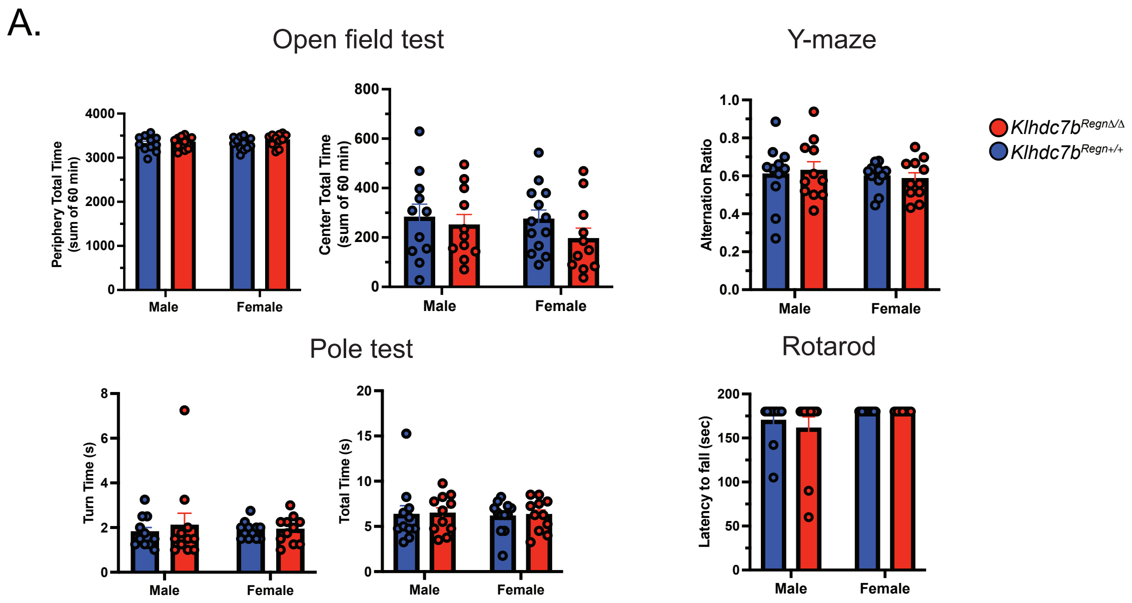


**Supplemental Figure 1. Behavioral testing in** *$Klhdc7b^{Regn+/+}$*, **and** *$Klhdc7b^{Regn\Delta/\Delta}$* **mice.** For the open field, mice were 8-12 weeks of age. For the Y-maze, pole test and rotorod, mice were 12-16 weeks of age. Two-way ANOVA was performed and found no significant main effect of genotype or sex on any of the behavioral tests performed. n = 10 mice per sex per genotype.

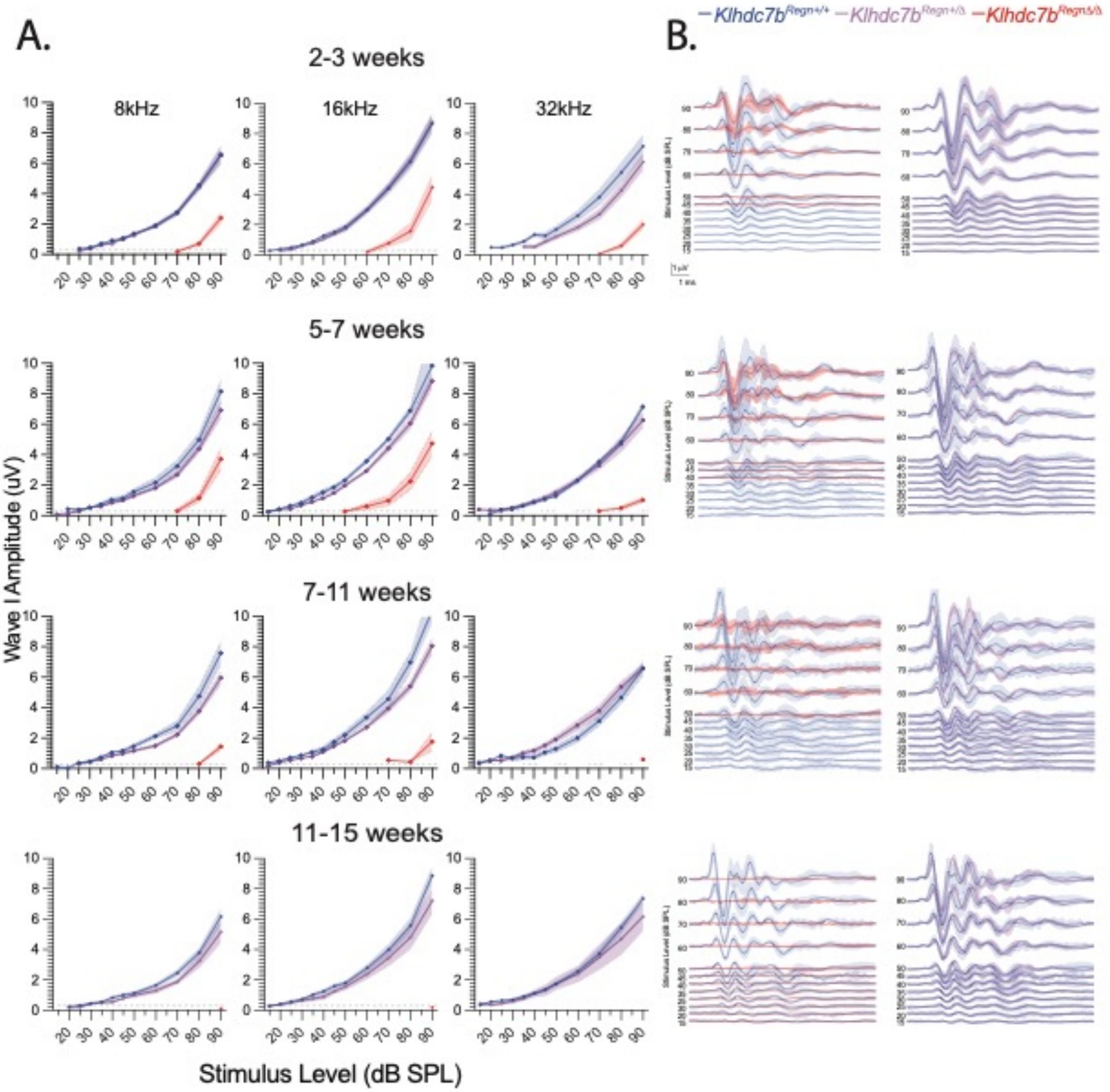
A.
B.
Klhdc7b Regn+/+
Klhdc7b Regn+/Δ
Klhdc7b RegnΔ/Δ
2-3 weeks
5-7 weeks
7-11 weeks
11-15 weeks
8kHz
16kHz
32kHz
Wave I Amplitude (uV)
Stimulus Level (dB SPL)

7

**Supplemental Figure 2.** Detailed time course recordings of $Klhdc7b^{Regn\Delta/\Delta}$, $Klhdc7b^{Regn+/\Delta}$, $Klhdc7b^{Regn+/+}$ mice at different time points. A. Wave I amplitudes for all time points. Analysis was performed via two-way ANOVA, which found a significant main effect of genotype. B. Average recordings at different time points. Left column, $Klhdc7b^{Regn\Delta/\Delta}$ and $Klhdc7b^{Regn+/+}$. Right column, $Klhdc7b^{Regn+/\Delta}$ and $Klhdc7b^{Regn+/+}$. Shaded area is standard error of the mean.

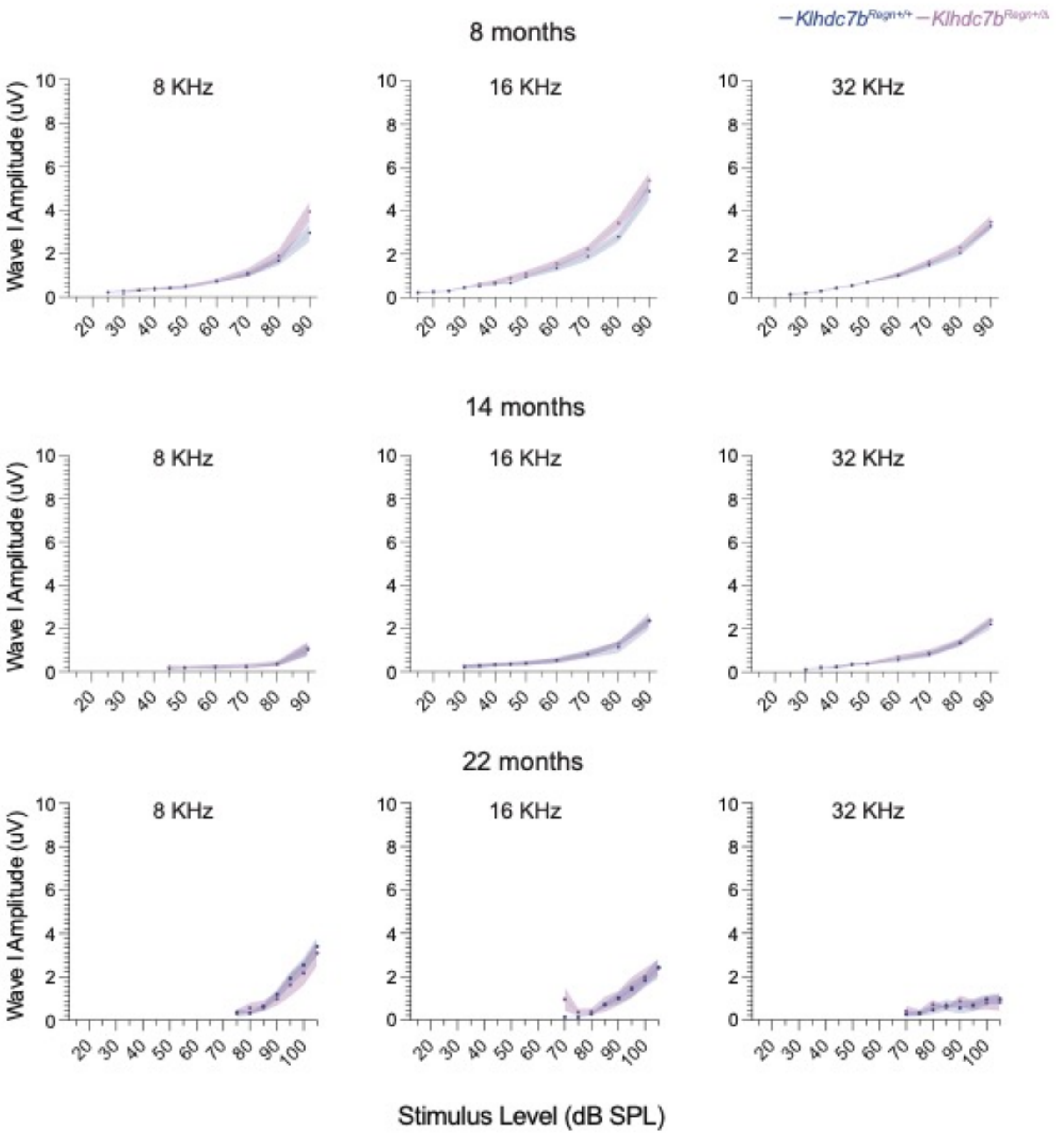

Klhdc7bRegn+/+
Klhdc7bRegn+/Δ
8 months
14 months
22 months
8 KHz
16 KHz
32 KHz
Wave I Amplitude (uV)
Stimulus Level (dB SPL)


13

**Supplemental Figure 3.** Wave I amplitudes from longitudinal recordings of *Klhdc7b*$^{Regn+/\Delta}$ and *Klhdc7b*$^{Regn+/+}$ mice at 8 months (top), 16 months (middle) and 22 months (bottom). Data was analyzed by two-way ANOVA and no significant differences between groups were found.

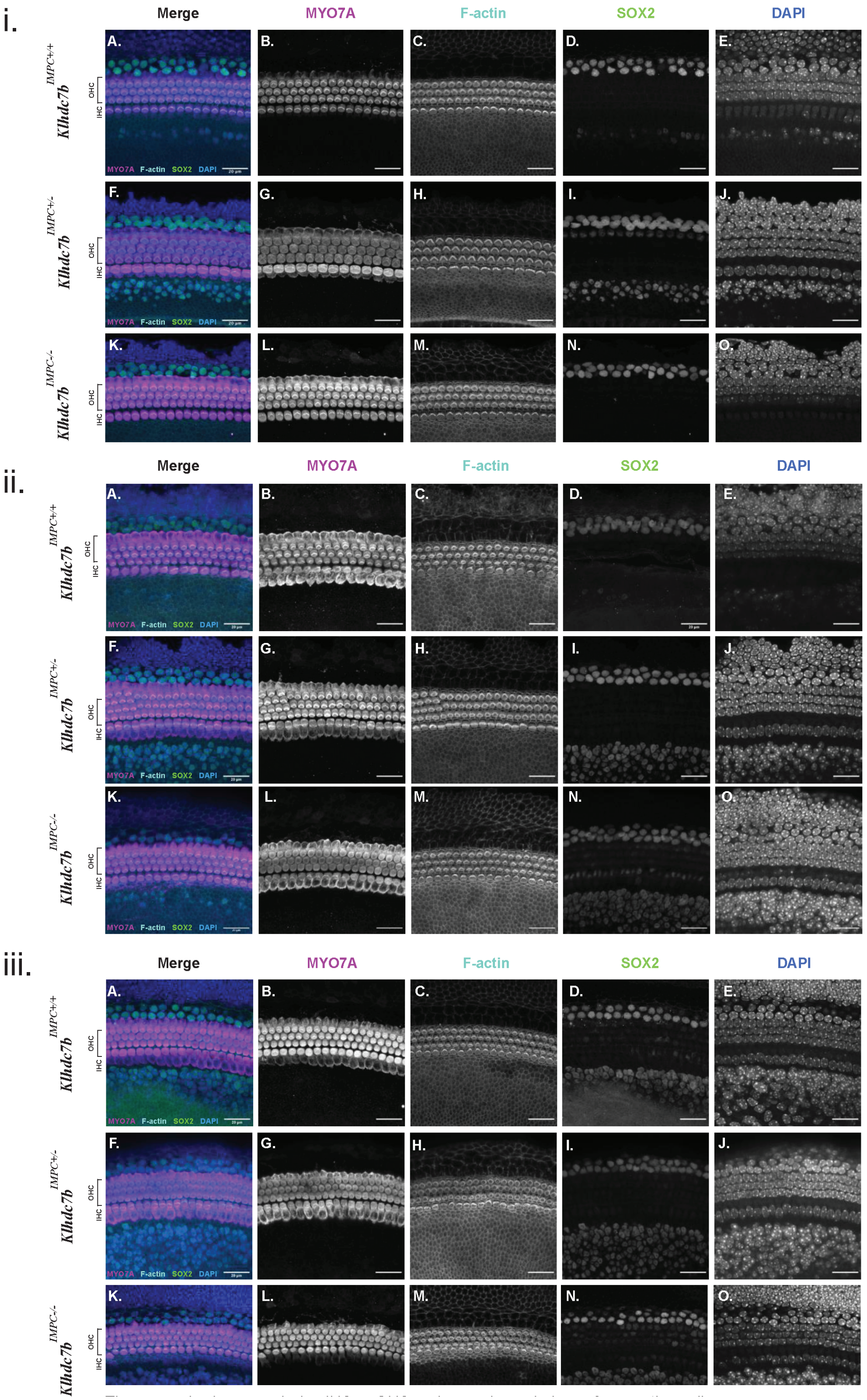

i.
ii.
iii.
Merge
MYO7A
F-actin
SOX2
DAPI
Klhdc7b IMPC+/+
Klhdc7b IMPC+/-
Klhdc7b IMPC-/-
OHC
IHC
A.
B.
C.
D.
E.
F.
G.
H.
I.
J.
K.
L.
M.
N.
O.

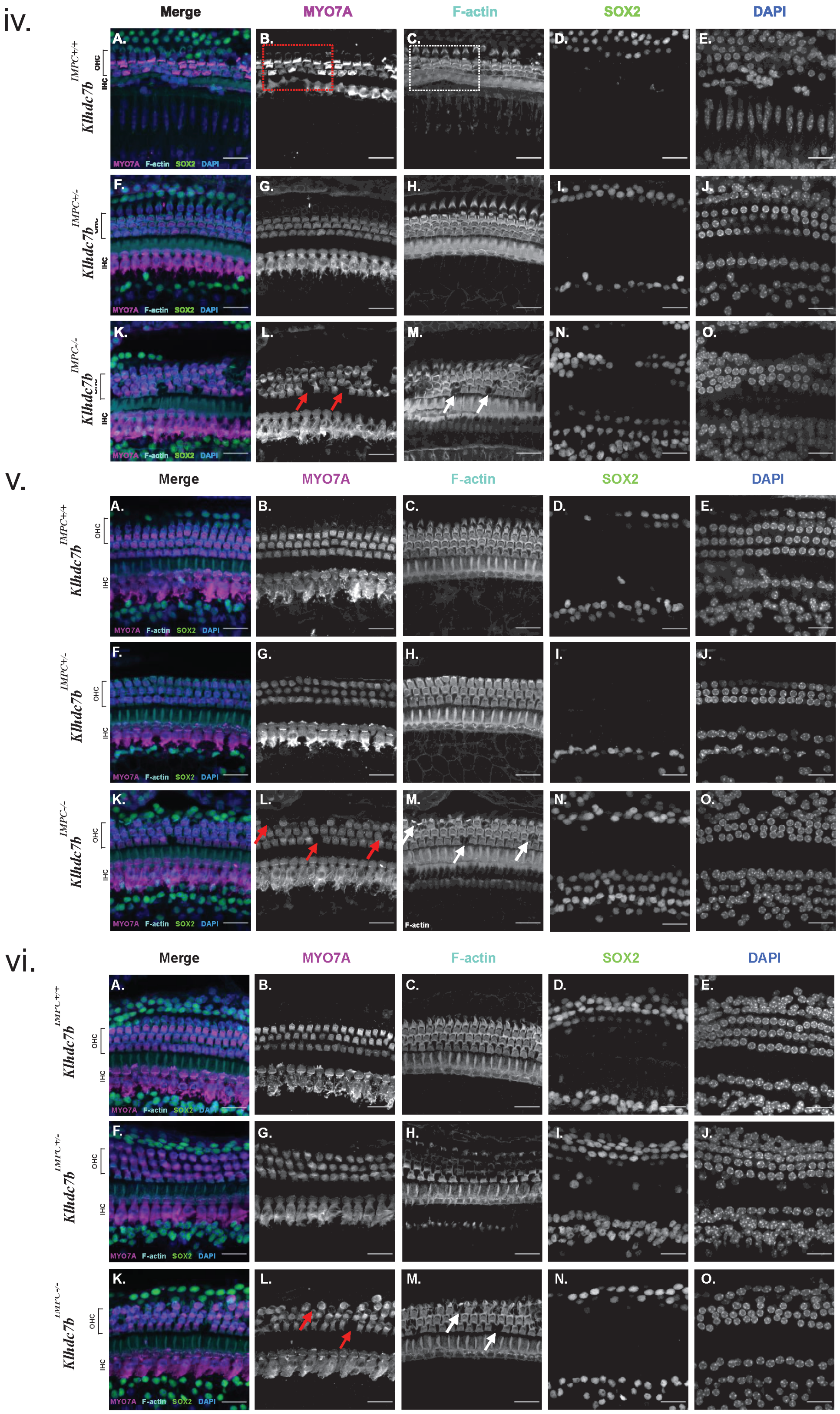
iv.
Merge
MYO7A
F-actin
SOX2
DAPI
Klhdc7b IMPC+/+
Klhdc7b IMPC+/-
Klhdc7b IMPC-/-
OHC
IHC
MYO7A F-actin SOX2 DAPI
v.
Merge
MYO7A
F-actin
SOX2
DAPI
Klhdc7b IMPC+/+
Klhdc7b IMPC+/-
Klhdc7b IMPC-/-
OHC
IHC
MYO7A F-actin SOX2 DAPI
F-actin
vi.
Merge
MYO7A
F-actin
SOX2
DAPI
Klhdc7b IMPC+/+
Klhdc7b IMPC+/-
Klhdc7b IMPC-/-
OHC
IHC
MYO7A F-actin SOX2 DAPI

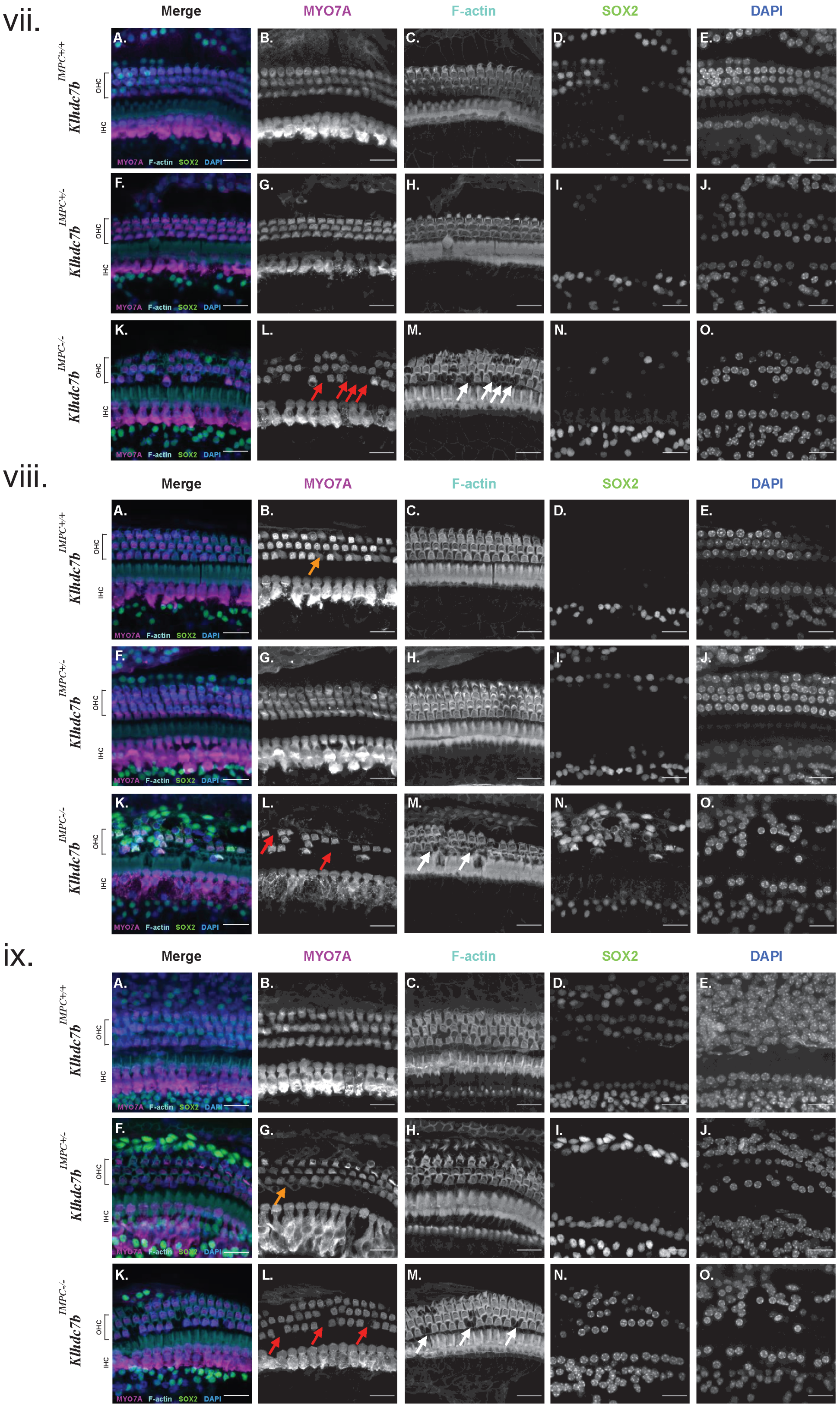
vii.
Merge
MYO7A
F-actin
SOX2
DAPI
Klhdc7b IMPC+/+
Klhdc7b IMPC+/-
Klhdc7b IMPC-/-
OHC
IHC
MYO7A F-actin SOX2 DAPI
viii.
ix.

**Supplemental Figure 4.** Cochlea whole mounts are shown in panels (i) to (ix) from the basal, mid and apical turn of P2 - P4 (i-iii), 3 week (P19-P21, iv-vi) and 8 week (P58-P60, vii-ix) mice in *Klhdc7b*$^{IMPC+/+}$, *Klhdc7b*$^{IMPC+/-}$, *Klhdc7b*$^{IMPC-/-}$ mice. Hair cells immunolabelled using Myosin VIIa (MYO7A) in magenta, actin cytoskeletal and stereocilia bundles labelled using Phalloidin (F-actin) in cyan, non-sensory epithelium immunolabelled using Sox2 in green, and nuclear immunostaining with DAPI in blue. OHC = Outer hair cells (indicating the three rows of OHCs). IHC = Inner hair cell. Scale bar = 20 µm, all images representative of $n \geq 3$ mice per genotype. The three rows of OHCs can be seen in parallel to the single row of IHCs (Myosin VIIa), on either side supporting cells are labelled using Sox2. The IHC and OHC are labelled at the side of panels A, F, K. Red and white arrows indicate examples of OHC loss. Some artefactual damage as part of mounting is present in viii and ix: orange arrows indicate where hair cells may appear to be missing but they are just folded over. Red and white box in iv also indicate some artefactual damage on mounting.

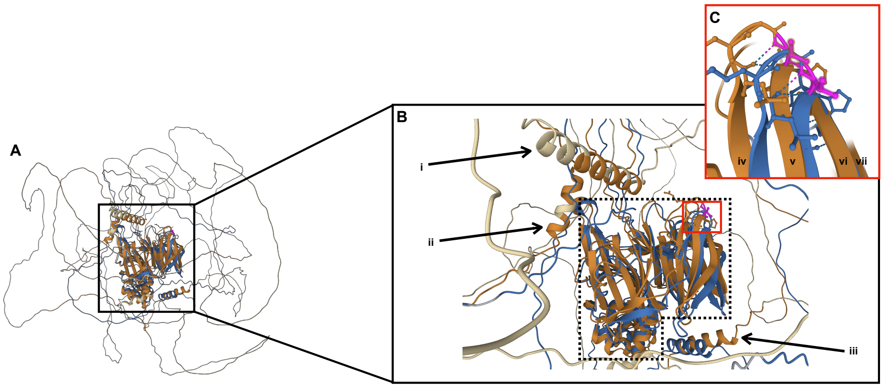


**Supplemental Figure 5.** Overlay analysis of the KLHDC7B long and short isoforms using the Pairwise structural alignment tool created by RCSB Protein Data Bank. **A.** Low resolution of both Klhdc7b isoforms, the long isoform (orange) and the short isoform (blue). The Kelch $\beta$-propeller of each variant is viewed in the same plane as the $\beta$-sheets. **B.** shows the KLHC7B proteins core at higher resolution and both the V1154 and V504 residues are annotated in magenta, **(i-ii & iii)** show $\boldsymbol{\alpha}$-helixes in close proximity with the Kelch $\beta$-propellers; however, i and ii are only within the long isoform structure and although iii is found in both isoforms, it can be seen that they do not appear to align completely. **C.** shows an enhanced resolution of the fifth Kelch motif in both isoforms, the valine residue is located within an inter-blade loop, and it can be seen than positions of the first two $\beta$-sheets of the long and short isoforms appear **(iv-v)** to be different but the third and fourth $\beta$-sheets $\beta$-propeller **(vi-vii)** in similar positions. This suggests that the Kelch $\beta$-propeller is subtly different between these two isoforms.

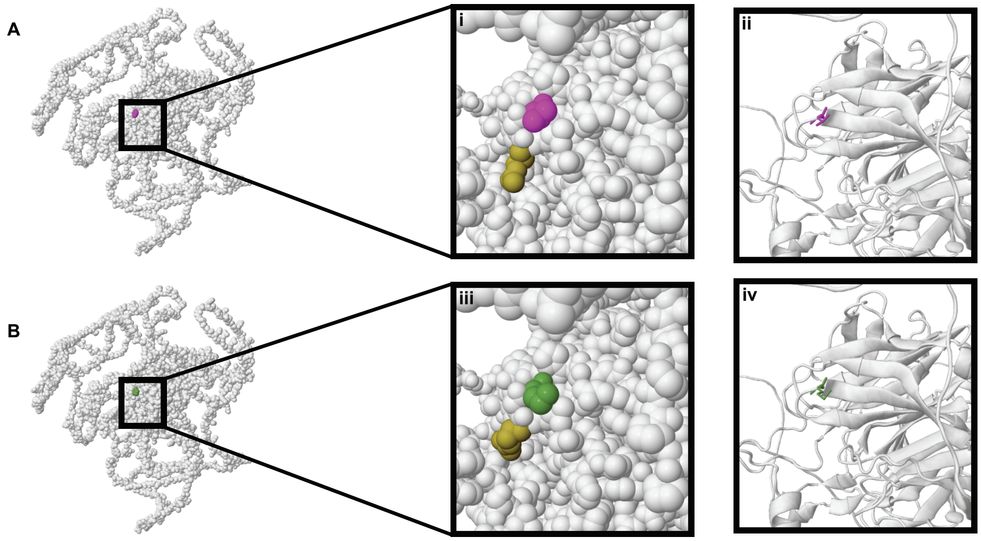


**Supplemental Figure 6.** Calotte space-filling modelling of Val and Met alleles on the long isoform of KLHDC7. Missense 3D modelling predicts the Val1154Met variant long is structurally damaging on the long isoform following contraction of cavity volume by 395.928 Å^3, the threshold for this criterion is an expansion or contraction of the cavity volume ≥70 Å^3 and it defines a cavity as either within the core regions of the protein or a pocket on the surface. **A.** KLHDC7B Val1154 (magenta) and **B.** KLHDC7B Met1154 (green). Boxes i and iii show the two residues at higher resolution and ii and iv show the same region at high resolution using as cartoon modelling. There is a subtle change in an adjacent space fill region (seen in yellow) following the missense mutation.

**Supplemental Table 1. VEP predictions for the effect of the ARHL associated Val>Met variant on the short and long form of Klhdc7b.**

| SNP ID | Ensembl Transcript ID | Amino Acid | Position | SIFT | PolyPhen |
|---|---|---|---|---|---|
| rs36062310 | ENST00000395676.4 | V/M | 504 | Tolerated (0.06) | benign (0.278) |
| rs36062310 | ENST00000648057.3 | V/M | 1154 | Deleterious (0) | possibly damaging (0.812) |
