## Supplemental Methods for "KLHDC7B, a novel gene associated with age-related hearing loss in humans, is required for the maintenance of hearing in mice"

**qPCR probes**

| Species | Name, gene | Probe | Forward primer | Reverse primer | company |
| --- | --- | --- | --- | --- | --- |
| mouse | L+S  mouse *Klhdc7b* | TTTATGCCATTG  GTGGCGAGTGC | GGTGGCCCTG  GATGGAATG | TCTGTGCGTG  GGTCATAGC | IDT (Integrated DNA technologies) |
| mouse | L mouse *Klhdc7b* | TCCCCATCATGAT  CCACTTTACCCC | CCGCAAAGCAA  AGGATATTCCTG | ATGCCTCCAG  CTGCATCTCTA | IDT |
| mouse | Mm01310009_  m1_Drosha,  Drosha |  |  |  | Thermofisher |

**Generation of *Klhdc7b^Regn^****^Δ^****^/^****^Δ^* **knockout mice**

*Summary description of sequences*

| **Description** | **Features** |
| --- | --- |
| Mus Musculus *Klhdc7b*, mRNA, NM_001160178 | Length: 5,764 bp  CDS: 859-4,650  Exons: 1-5,764; |
| Mus musculus Klhdc7b, protein | Length: 1,263 aa |
| LacZ replacing *Klhdc7b,* DNA | Length: 3,075 bp |
| LacZ replacing *Klhdc7b,* protein | Length: 1,026 aa |
| 5’ mouse UTR // **Start**, **Acc65** // 5’ LacZ | 5’mouse // 5’ LacZ |
| 3’ LacZ //**Stop** // (*LoxP*) // **NheI** // 3’ mouse | 3’ LacZ, 3’ mouse UTR |
| Primers and probes for loss of allele and gain of allele assays for *Klhdc7b* KO | Table 1 |

*Table 1*

| **NAME** | **Primer** | **Sequence (5’-3’)** |
| --- | --- | --- |
| 4929mTU | Forward  Probe (BHQ)  Reverse | TGGGAGGCGGTTGTACTC  ACCTTGCTGGCACTAGCGGTGGT  GCAAAGCCAGTGCTAAGGT |
| 4929mTD | Forward  Probe (BHQ)  Reverse | GGCTTCCTATACCGCTTTGAC  TCCAGCTATCGCACTTATCTGA  CTCCAGGAGCCGGTCACT |
| LacZ | Forward | GGAGTGCGATCTTCCTGAGG |
|  | Probe (BHQ) | CGATACTGTCGTCGTCCCCTCAAACTG |
|  | Reverse | CGCATCGTAACCGTGCATC |
| Neo | Forward | GGTGGAGAGGCTATTCGGC |
|  | Probe (BHQ) | TGGGCACAACAGACAATCGGCTG |
|  | Reverse | GAACACGGCGGCATCAG |

*Description of BAC clone generation for* *Klhdc7b^RegnΔ/Δ^ mice*

A BAC clone RP23-241G24 containing a mouse *Klhdc7b* gene was used and modified as follows. Briefly, a DNA fragment was generated to include a mouse 5’ homology nucleotide sequence of 100 bp (mHU), a LacZ gene (3,075bp) downstream and in frame with the ATG starting site of *Klhdc7b* gene, followed by a self-deleting neomycin cassette of 4,809 bp, and a 3’ mouse homology sequence of 100 bp (mHD). This DNA fragment was used to modify BAC clone RP23-241G24 through homologous recombination in bacterial cells. As result, a full KO of the region encoding mouse *Klhdc7b* genomic fragment of 3,787 bp in the BAC clone was replaced by the LacZ, a Neo-self-deleting cassette (SDC) of 8,202 bp. The entire mouse *Klhdc7b* orf (mm9, chr15:89,215,351-89,219,137) was replaced with the LacZ-Neo SDC leaving intact 5’ and 3’ UTRs. The resulting modified BAC clone included, from 5’ to 3’, (i) a 5’ mouse homology arm containing about 140.5 kb of mouse genomic DNA including a mouse *Klhdc7b* 5’ UTR and ATG; LacZ cDNA of 3,075 bp, a self-deleting Neomycin cassette of about 4,809 bp, followed by a 3’ mouse homology arm of 12.6 kb containing the mouse *Klhdc7b* 3’ UTR and the remaining mouse genomic DNA in the original BAC clone.

**Primers for Klhdc7b^IMPC-/-^ mouse genotyping**

| **Primer name** | **Sequence 5' → 3'** | **Primer Type** |
| --- | --- | --- |
| Klhdc7b-DEL1258-UNI_F | CCTCTGAAGGCGCCATCTTC | Universal Forward |
| Klhdc7b-DEL1258-WT_R | TGCTTGGTTTCATCCCCTCTC | Wildtype Reverse |
| Klhdc7b-DEL1258-MUT_R | CCTAGCGCCACCATTCACTG | Mutant Reverse |

**Antibodies and Stains**

| **Antibody Name (target)** | **Source** | **Catalog** | **primary/**  **secondary** | **host species** | **dilution** |
| --- | --- | --- | --- | --- | --- |
| Myo7A | proteus | 25-6790 | primary | rabbit | 1 in 1000 |
| KLHDC7B Immunogen: entire short isoform of human protein | custom | Custom | primary | rabbit | 1 in 200- 600 |
| Sox2 | BDPharmingen | AB_1645334 | primary | mouse |  |
| DAPI | Thermofisher | 62248 | cell stain | N/A | 1 in 1000 |
| Alexa Fluor 647 phalloidin | Thermofisher | A22287 | cell stain | N/A | 1 in 500 |
| phalloidin-Atto 647N | Sigma | # 65906 | cell stain | N/A | 10nM |
| donkey anti-rabbit Alexa 488 | Thermofisher | A32790 | secondary | donkey | 1 in 1000 |
| donkey anti-mouse Alexa 568 | Thermofisher | A10037 | secondary | donkey | 1 in 1000 |
| goat anti-rabbit (IgG H+L) Alexa 488 | Thermofisher | A-11008 | secondary | goat | 1 in 1000 |
| goat anti-(mouse IgG2a) Alexa 546 | Thermofisher | A-21133 | secondary | goat | 1 in 1000 |

**RNA scope reagents**

| **Reagent** | **Source** | **Catalog/reference #** |
| --- | --- | --- |
| Mouse *Klhdc7b* long probe only | ACD bio | 1088961-C1 |
| Mouse *Klhdc7b* overlapping probe | ACD bio | 1136591-C2 |
| Mouse positive control probes | ACD bio | 320881 |
| negative control probes | ACD bio | 320871 |
| protease and peroxide kit | ACD bio | 322381 |
| fluorescence detection kit | ACD bio | 323110 |
| Opal 520 | Akoya biosciences | OP-001001 |
| Opal 570 | Akoya biosciences | OP-001003 |
| Anti-MYO7A antibody | Proteus | 25-6790 |
| Donkey anti-rabbit Alexa 647 | Thermofisher | A31573 |
| Prolong gold | Thermofisher scientific | P36934 |

**Behavioral testing**

One week before starting behavioral evaluations mice were handled for five minutes over two consecutive days to reduce the impact of handling on behavioral readouts. On experiment day, mice were additionally acclimatized to the experimental room at least for an hour.

*Open Field.* Mice were placed in an open field Plexiglas arena (40.6 cm x 40.6 cm x 38.1 cm) containing two horizontal laser beam detection arrays. The tracking software Motor Monitor (Kinder Scientific) for Windows (Microsoft) was used to measure movement in X and Y axis. Groups of eight animals, counterbalanced across experimental conditions, were tested concurrently for 60 minutes. Behavior was measured in five-minute intervals and a total count for each measure for the 60 minutes was calculated by summing the twelve five-minute bins. Computed measures include: basic movements (any horizontal beam cross), immobility time (lack of horizontal and vertical beam crosses), fine movements (changes in body position not meeting criteria for ambulation, includes grooming and head movements), X+Y axis ambulation (complete relocation of the animal’s body), rears (vertical beam crosses), rearing time (time spent breaking vertical beams), rest time (lack of beam crosses lasting longer than 15 seconds), and total distance traveled (computed from known distances between beams and total beam crosses).

*Rotorod.* Animals were trained to walk on the Rotorod equipment (IITC Life science) containing 5 separate lanes with individuals rotating drums. Training consisted of 3 separate runs of 180 seconds where rotation speed was progressively increased: trial 1 (0 to 15 rpm, rotations per minute), trial 2 (7 to 15 rpm) and trial 3 (7 to 15 rpm). Mice were replaced back onto drum if they fell off. An animal was considered ready for testing phase if able to walk on the drum for at least 150 seconds. Three testing trials per animal were run (inter-trial interval = 30 minutes). During testing trials rotation speed was 15 rpm and mice were assessed for 180 seconds until a fall occurred. The median latency to fall was computed for each mouse.

*Pole Test.* The pole test was carried on a metal rod (50 cm long, 1 cm diameter) mounted on a square base that was buried under home cage flooring. During adaptation trials each mouse was placed three times on the rod (head down) to facilitate learning of pole-descent. In testing trials mice were placed on the top part of the rod (head up) and evaluated for the ability to turn around and descend the pole safely. The time to make a full 180 degrees turn and latency to reach the floor was recorded for 5 consecutive times. Trials where mice fell off the rod were annotated with the maximum trial duration (60 seconds). Average time to turn around and descend were calculated for the best 4 (out of 5) consecutive trials.

*Y-maze (spontaneous alternation).* Mice were placed in a Y-maze apparatus constituted of three plastic arms (14.5 cm × 3.5 cm × 13.5 cm) separated by a 120° angle from each other. At the beginning of each trial mice were positioned at the center of the maze and allowed to explore for 8 minutes. Using Ethovision system (Noldus, The Netherlands) body position was tracked over time to compute the number of spontaneous arm alternations (visits to an arm that was previously not visited) and the maximum number of possible alternations after a given number of visits to a new arm. Alternation ratio (AR) was calculated by dividing spontaneous arm alternations over maximum number of possible alternations.

**ABR**

For recordings at Regeneron, equipment was calibrated each day using a microphone (model PCB 378C01, PCB Piezotronics, NY) placed at the same distance from the speaker as the mouse ear being recorded. Animals were anesthetized with an intraperitoneal injection of ketamine/xylazine (12 mg/kg, 0.5 mg/kg) and placed in a heated cage. Puralube ointment was placed on the eyes after several minutes once the animal was no longer responsive.

Once the mouse was fully anesthetized it was placed on a Gaymar heating pad (Gaymar Industries, NY) in the sound booth. Electrodes were plugged into a Medusa 4Z preamplifier (Tucker-Davis Technologies, FL). Three lead, 13mm needle electrodes were placed subdermally with the lead electrode at the cheek of the animal (near the cochlea), the reference electrode at the midline of the skull on top of the head, and the ground electrode in the contralateral cheek. The ear being recorded was positioned 7.5 cm away from the speaker, in an open field configuration. After recordings, animals were placed in a heated recovery cage and returned to the home cage once ambulatory.

For recordings at Harwell, mice were anesthetised by intraperitoneal injection of ketamine (100 mg ml^−1^ at 10% v/v) and xylazine (20 mg ml^−1^ at 5% v/v) administered at the rate of 0.1 ml per 10 g body mass. Animals were placed on a heated mat inside a sound-attenuated chamber (ETS Lindgren) and electrodes were placed subdermally over the vertex (active), along the right mastoid (reference) and on the left flank (ground). Animals were recovered using 0.1-0.2 ml of anaesthetic reversal agent atipamezole (5 mg ml^−1^ at 1% v/v).

**ABR analysis**

For automated threshold calling, the covariance between pairs of adjacent decibel traces was calculated and plotted. These points were fit to a curve using a sigmoid or logarithmic function, and the decibel level at which the function crossed below a set criterion level was recorded as the threshold of hearing. The threshold values called by the algorithm were compared to manually called thresholds. Traces with no discernible ABR response were called at 100 dB. If the difference between the manual and automated thresholds was greater than 15 dB, traces were examined and the manual threshold was used; otherwise, the automated threshold was used.

For wave 1 amplitude and latency, peaks were detected using a semi-automated method where a peak and trough was estimated within a time window to encompass wave 1, then checked by a user and corrected if necessary.

**3D microscopy**

*Staining and clearing protocol.* Cochlea were collected, stored in PBS, and decalcified with immunocal overnight, then rinsed 3x with PBS at room temperature. All washes and incubations were conducted with shaking. The decalcified bone was carefully cut around the spiral to improve reagent penetration. Cochleae were washed 3x 2 hours with PBS at room temperature, then blocked for two hours at 37°C, using standard blocking solution as for immunohistochemistry. MYO7A antibody was diluted in blocking solution at 1:200-1:300, then samples were incubated in primary antibody for 48-72 hours at 37°C. Samples were washed 3x for two hours in PBS at room temperature. Secondary antibody (Donkey anti-rabbit Alexa 568, thermofisher, A10042) and a nuclear stain (Sytox Deep Red, Thermofisher, S11381) were diluted at 1:200 and 1:400, respectively, in blocking solution. Samples were incubated in this solution overnight at 37°C, then washed 3x for two hours in PBS at room temperature. Samples were incubated at room temperature in a dilution series of concentrations of methanol (30% methanol, 50%, 70%, 90% and 98%) plus 2% Tween-20 in ddH_2_O for 8-18 hours each. Samples were then placed in a glass jar and incubated at room temperature in 100% Ethyl Cinnamate (ECi) for two hours, then placed in fresh ECi and incubated at room temperature for ~72 hours. Samples were stored in ECi at room temperature or at 4°C until imaging.

*Imaging.* Samples were imaged using an UltraMicroscope Blaze (Miltenyi Biotec). The 12x objective was used with the 1.67x tube lens for a total magnification of 20x with 100% beam width and beam thickness of 3.9um, with illumination from both sides and five steps of dynamic focus. Z stacks were spaced at 2 μm. Samples were imaged using the 561 and 640 lasers for Alexa 568 and Sytox deep red, respectively.

**Automated Hair cell counting in intact cochlea**

*Virtual Dissection*. After imaging, files were processed and converted to Imaris (v10.1.0) files. To minimize the need for large computational resources, images were cropped in 3D to focus solely on the hair cells. Hair cells were virtually dissected using the ‘filament’ tool in Imaris by tracing the inner hair cells from the apex to the base as described by Hutson et al. (27). The filament diameter was set to 120 mm to capture both inner and outer hair cells. To create a mask, the Imaris ‘Filament to Channel’ Xtension was used to generate a new channel of data based on the filament. The channel was converted using the ‘Surface’ tool in Imaris based on the signal from the new filament channel. The generated surface was used to mask the original data channels containing both nuclear signal and hair cell fluorescence. Data from both masked channels were exported as TIFF images (along the *z*-axis) for automatic counting of cells. In addition to masking the combined signal from the inner and outer hair cells, an identical procedure was employed to isolate signal from only the inner hair cells. All subsequent analysis was performed using Python (v3.9.6) on a 32-core cloud-based compute cluster with 128 GB of RAM.

*Automated Hair Cell Counting*. Masked hair cell nuclear and fluorescence images were tiled into 150 x 150 pixel^2^ sub-images to further maximize processing on data in the limits of available memory during automated counting (I_nuc_ , I_FLR_ ,respectively). Simultaneously, an outer hair cell mask (M_OHC_) was generated using the inner hair cell mask (M_IHC_) by:

1. Smoothing each *z*-slice image via morphological closing
2. Radially offsetting the boundaries of inner hair cell mask by 100 pixels from the center of the cochlea spiral.
3. Morphological dilation of the offset mask by a disk (radius = 45 pixels)

For all further analysis, both the inner and outer hair cell masks were tiled in the same manner as the original nuclear and fluorescence images. To isolate hair cells, the masks for inner and outer hair cells were multiplied against the tiled nuclear and fluorescence images to extract pixels within each image that corresponded to either cell type (M_IHC_ x I, M_OHC_ x I). Hair cell fluorescence images were thresholded to exclude background fluorescence (I_FLR, thresh_). After excluding background fluorescence, we calculated the standard deviation per *z*-slice in each tile, and used local maxima per tile to identify *z*-slices of interest, including additional *z*-slices defined as z-slices within the full-width-at-half-maximum (FWHM) of the local maxima peaks. The identified *z*-slices were then processed individually to extract the hair cells using a hair cell selection function.

Briefly, a multi-level (four level) Otsu threshold was used to generate an image with four discrete pixel values from which the highest-valued pixels were isolated for both the nuclear and the fluorescence images per *z*-slice. A four level Otsu threshold was selected based on the distribution of signal in both the nuclear and fluorescence images. As the hair cell fluorescence and nuclear signal overlapped maximally at the center of the hair cell, it was assumed that the product would isolate hair cell signals. Therefore, the product of the maximal intensity value was calculated to extract the hair cell signal (I_nuc_ x I_FLR, thresh_). To further remove contribution of signal from nearby cochlear supporting cells that overlapped with the hair cell fluorescence, we removed smaller objects using an area threshold (<50 pixels^2^ or ~ 5.3 mm^2^).

To generate an individual hair cell mask, we calculated the distance between each pixel in a cell and the background. We then isolated the location of the maximum distance per cell (at the center of the cell) and marked the center of each hair cell. At each location (cell center), a uniform circle was drawn around the cell nucleus and used to label hair cell containing pixels in the hair cell signal images. To label and mask each individual cells in the *z-*slice, we applied a watershed algorithm to this image. At each labelled cell’s centroid, the intensity was recorded for individual cell mask generation. A circle was then drawn from the centroid with a radius equal to the distance at which the signal per each cell decreased below 95% of the previously recorded centroid intensity. To ensure that cells were not counted more than once, each cell was compared to detected cell objects in the prior 10 slices and overlapping cell objects were removed. For each tile, we multiplied the labelled cell objects with the original masked cell image (I_nuc_ x I_FLR, thresh_). This process was performed for each cell type (IHC and OHC). Once all tiles had been processed, all tiles were stitched back together to create a final XYZ stack (I_IHC_, I_OHC_, respectively).

All processing up to this point used 3D datasets, ensuring the preservation of positional relationships of all hair cells in three-dimensional space. To accurately count the hair cells from apex to base, a custom processing pipeline was developed to unwrap the cochlea in a sequence that reflects its natural configuration. Despite meticulous efforts used during tissue embedding, the alignment of cochleae with the global axis was not perfect. To compensate for this, each *z*-slice was analyzed sequentially, with cells object being counted as they became visible and positionally stored in a growing map. When newly-visualized cell objects appeared more than 100 pixels from the existing cell object, or when cell objects appeared to be unwrapping in two directions, a new map was initiated to track the “far” or “diverging” cell objects.

After mapping the entire cochlea in 3D, the center of the maximum intensity projection (MIP) image (along the *z-*axis) was used to select one of the maps created above as the starting point to unwrap the cochlea. To select the starting map, we calculated the minimum distance between the center of the MIP and the *(x,y)* positional coordinates of all the maps we had generated. Once the start map (L_1_) was identified, the map was traced to its opposite end, and the rotational direction from the center of the MIP was identified (either clockwise or counterclockwise) based on the change in the angle of rotation from the start to the end of the map. Subsequently, the remaining maps were examined to determine if the starting map (L_1_) was next to another map in 3D space or was connected to either the base or apex of the cochlea. If a nearby map was identified near the terminal end of L_1_, the nearest point of this new map was linked to the L_1_ map. This process was repeated until the final map reached either the apex or base of the cochlea.

After unwrapping part of the cochlea, the script would return to the start map of L_1_ (middle of the cochlea) and reverse the rotational direction. It then searched for the nearest map to the starting point in 3D space, tracing the maps until reaching the opposite end of the cochlea (either apex or base). Upon completion of this process, the shorter of the two linked maps (assumed to contain the apex due to its length) was flipped such that the last map added became the starting point and the start map of L_1_ was the end point. This map was then merged with the longer of the two linked maps (assumed to contain the base). This resulted in a naturally ordered mapping of the cochlea from the apex to the base. Simultaneously, accumulation of both inner and outer hair cells counts were traced along the finalized mapping, and cochlea length was calculated. Subsequently, a tonotopic map of the cochlea was constructed using the outer hair cell length from the apex to the base (25). Tonotopic mapping was calculated using:

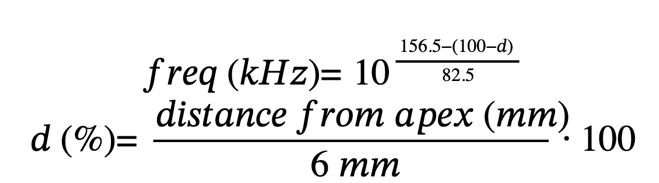

The normalization factor of 6 mm was selected based on the reported average length of a mouse cochlea from apex to base (28).

Results for all 12 cochleae were aggregated by excluding areas with damage to hair cells before assessing the rate of change in inner and outer hair cell counts. As the counting of cells was automated, individual IHCs were not mapped to corresponding rows of OHCs making instantaneous assessment of OHC to IHC ratio challenging. To account for this, we instead assessed the rate of change in OHC count and IHC count over discrete bins of cochlea length to extract an averaged ratio of OHC to IHC counts. The OHC to IHC ratio was calculated using 1 mm bins with ranges defined as 0 to 1 mm, 1.01 to 2 mm, 2.01 to 3 mm, 3.01 to 4 mm, and 4.01 to 5 mm.  Through the use of binning and tracking rate of change in counts of each cell, removed the need to map each IHC to a corresponding row of OHCs while still providing a meaningful assessment of OHC to IHC ratio.
